# Functional development of eye movements and visuomotor circuits in lampreys

**DOI:** 10.1101/2023.09.06.556551

**Authors:** Marta Barandela, Carmen Núñez-González, Daichi G. Suzuki, Cecilia Jiménez-López, Manuel A. Pombal, Juan Pérez-Fernández

## Abstract

Animals constantly redirect their gaze away or towards relevant targets and, besides these goal-oriented responses, stabilizing movements clamp the visual scene avoiding image blurring. The vestibulo-ocular (VOR) and the optokinetic reflexes are the main contributors to gaze stabilization, whereas the optic tectum integrates multisensory information and generates orienting/evasive gaze movements in all vertebrates. Lampreys show a unique stepwise development of the visual system whose understanding provides important insights into the evolution and development of vertebrate vision. Although the developmental emergence of the visual components, and the retinofugal pathways have been described, the functional development of the visual system and the development of the downstream pathways controlling gaze are still unknown. Here, we show that VOR followed by light-evoked eye movements are the first to appear already in larvae, despite their burrowed lifestyle. However, the circuits controlling goal-oriented responses emerge later, in larvae in nonparasitic lampreys but during late metamorphosis in parasitic lampreys. The appearance of stabilizing responses earlier than goal-oriented likely reflects their evolution, and its stepwise emergence offers a unique opportunity to isolate the functioning of their underlying circuits.

## Introduction

Ocular movements have gradually emerged across evolution depending on new motor and perceptual needs. The oldest eye movements appeared with the aim of immobilizing the visual image on the retina: the vestibulo-ocular reflex (VOR), which maintains clear vision by ensuring stabilization of the eyes during head movements, and the optokinetic reflex (OKR), that keeps the visual field clamped on the retina when it is shifted (Land, 2015). These reflexes have been maintained throughout evolution and their appearance dates back in the phylogenetic tree of vertebrates to the position of the lamprey, which belongs to the oldest group of living vertebrates (Rovainen, 1976; Wibble et al., 2022). The interface for the VOR lies in the vestibular nuclei, divided in lampreys in the anterior (AON), intermediate (ION) and posterior (PON) octavomotor nuclei, which receive vestibular information from the labyrinth and project to the three motor nuclei that control the extraocular muscles: oculomotor (nIII), trochlear (nIV) and abducens (nVI; Pombal et al., 1996; Fritzsch, 1998). As in other vertebrates, the pretectum (PT) generates OKR, sending visual information to the oculomotor nuclei both directly and through the vestibular nuclei (Giolli et al., 2006; Masseck and Hoffmann, 2009; Wibble et al., 2022). Thus, the circuits controlling these reflexes are very similar to other vertebrates meaning that they appeared at the dawn of vertebrate evolution and are largely conserved.

Apart from gaze stabilization, lampreys also possess the main centers involved in goal-oriented gaze redirection. As in mammals, the optic tectum (OT; its mammalian counterpart being the superior colliculus) has a layered structure and receives visual information that forms a retinotopic map of the surrounding space, aligned with other sensory modalities (de Arriba and Pombal, 2007; Jones et al., 2009; Kardamakis et al., 2016). Tectal outputs to premotor areas in the brainstem form motor maps aligned with the sensory maps, and sensory information is integrated by the tectal circuits that can prioritize the salient stimuli and encode the appropriate behavioral response by generating orienting or evasive movements (Saitoh et al., 2007; Kardamakis et al., 2015; 2016; 2017; Suzuki et al., 2019). The OT also receives other inputs that can modulate its motor commands, including those from the basal ganglia and cortex (Stephenson-Jones et al., 2011; Ocaña et al., 2015; Pérez-Fernández et al., 2014; 2017; 2021). Additionally, a visual area with retinotopic representation is present in the lamprey pallium, suggesting that a primordial blueprint of the visual cortex is present in these animals (Suryanarayana et al., 2020), and therefore all the main visuomotor areas are present in lampreys (Suryanarayana et al., 2021).

The visual system of lampreys shows a unique development, which has been suggested to reflect its evolution, because of their complex life cycle (Suzuki and Grillner, 2018). As larvae, lampreys are photophobic, filter feeders, and their eyes are immature in form of eyespots covered with skin, presenting a flattened lens and thus not being functional to form images (Kleerekoper, 1972). After a very long larval period (lasting for five to seven years; Hardisty, 2006), lampreys undergo a metamorphosis in which some species develop structures for their new parasitic life that include functional eyes (Kennedy and Rubinson, 1977; Suzuki and Grillner, 2018). The structures and brain circuits necessary to use the visual system for advanced behaviors develop stepwise during the larval period, and the visual system completes its development during metamorphosis (Kennedy and Rubinson, 1977; Suzuki and Grillner, 2018). In early larvae (< 60-70 mm), retinopretectal connections are already established, and direct retinal projections to the nucleus of the medial longitudinal fascicle (nMLF), which projects to the brainstem and spinal cord (SC), generate a simple circuit possibly involved in negative phototaxis (de Miguel et al., 1990; Cornide-Petronio et al., 2011; Suzuki et al., 2015a, b; Suzuki and Grillner, 2018). At this stage, the OT shows an immature state and has no input from the retina, which is formed only in larvae larger than 70 mm (de Miguel et al., 1990; Cornide-Petronio et al., 2011). The OT completes its development during metamorphosis, expanding and completing its lamination. The extraocular muscles also develop gradually during the larval period, starting before the development of the retinotectal projection (Suzuki et al., 2016), and it appears that the oculomotor, trochlear and abducens neurons that innervate those muscles also develop gradually, but their differentiation is earlier than that of the extraocular muscles (Fritzsch et al., 1990; Pombal et al. 1994; Suzuki and Grillner, 2018). Retinal development is also gradual. Prolarvae and early larvae exhibit a primary retina without horizontal and amacrine cells, essential for image-forming vision (Villar-Cerviño et al., 2006), and during the larval period the retina expands and undergoes extensive cell differentiation completing its development during metamorphosis (Kennedy and Rubinson, 1977). During this period, the eye also acquires a chambered structure with a spheric lens, and the overlying skin becomes transparent, becoming truly functional (Dickson and Collard, 1979). Regarding stabilizing gaze movements, the circuits mediating VOR are established during the larval period (Pombal et al., 1996; Fritzsch, 1998), although whether this reflex is functional at this stage is unknown. Regarding OKR, no data exist in larva.

The large degree of similarities between the visual system of lampreys and that of other vertebrates, together with its gradual development through a very long period, offer a unique window to isolate functional developmental aspects of the different visuomotor behaviors and their underlying circuits (Suzuki and Grillner, 2018). Although the order of appearance of the different visual components, and the development of the retinofugal pathways have been described (Kennedy and Rubinson, 1977; de Miguel et al., 1990; Cornide-Petronio et al., 2011), their functionality is unknown. Moreover, although some of the motor pathways that generate orienting, evasive, and eye movements are well described in adults (Saitoh et al., 2007; Kardamakis et al., 2015; 2016; 2017; Suzuki et al., 2019), their development is still unknown. In this study, we show that larval lampreys have coordinated eye movements in the form of VOR, and that eye movements can also be evoked by light stimuli. However, the direct pretectal and tectal motor outputs to reticulospinal neurons are mostly established during metamorphosis, although these structures can trigger motor responses before, which are conveyed through polysynaptic pathways. Interestingly, tectal and pretectal projections are established earlier in non-parasitic lampreys. Given the high degree of conservation of the analyzed circuits, our results provide insights to understand the functional development of the vertebrate visual system.

## Results

### Larval lampreys exhibit coordinated eye movements

Given that the eye muscles already develop during the larval period (⁓60 mm length), and the innervation from the motor nuclei appears even earlier (Fritzsch et al., 1990; Pombal et al., 1994; Suzuki et al., 2016), we first explored whether this motor infrastructure is already functional in lamprey larvae even though they spend most of their life burrowed, and that their eyes are not fully developed and covered with skin. For this, we used a preparation exposing the eyes and the brain, so that we could stimulate different brain areas and monitor eye movements via video recordings and/or electromyograms (EMGs) in the extraocular muscles (Fig. 1A). The progressive development of the eyes during the larval period, in which they grow and protrude from the cartilage around the brain, can be seen in the preparations from larvae of different sizes (Fig. 1Bi-iii). In small larvae (Fig. 1Bi), the eyes appear immature and noticeably small. As the larva grows, the eyes enlarge (Fig. 1Bii), and this process continues during metamorphosis. It is only after metamorphosis that their eyes are fully developed (Fig. 1Biii; Kennedy and Rubinson, 1977; Suzuki and Grillner, 2018).

**Fig. 1.**
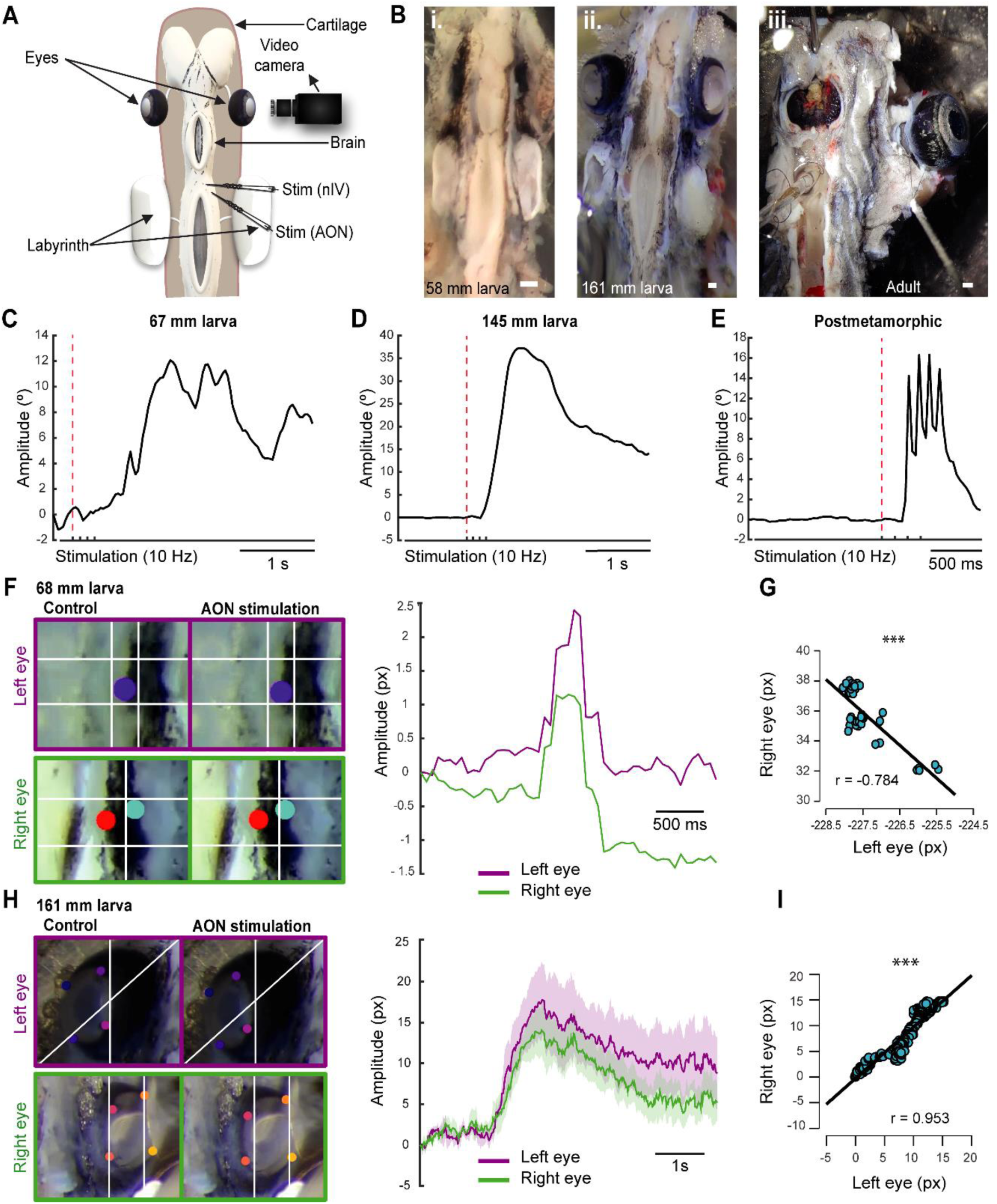
Lamprey larvae exhibit eye movements. (A) Schematic showing the *ex vivo* preparation used to monitor eye movements. (B) Preparations of larvae of 58 (Bi) and 161 (Bii) mm long showing eye development in relation to body size. A preparation of an adult animal is shown in Biii. (C-E) Traces representing the eye position in response to a four pulses stimulation (10 Hz) of the Anterior Octavomotor Nucleus (AON) in a 67 (C) and a 145 (D) mm larva, and a postmetamorphic (E) lamprey. The red dotted line indicates when the first pulse was applied. (F) Right: Traces showing the position of the right (green) and left (purple) eyes in response to a four pulses 10 Hz stimulation of the AON in a 68 mm larva, in which coordinated movements can be observed. Left: Frames showing the position of the eyes (top, left eye; bottom, right eye) before (left) and after (right) stimulation in the AON. Colored dots are the labels used to track eye movements. (G) Graph showing the correlation between the movements of the right and the left eyes of the representative 68 mm larva shown in F, indicating their coordination after stimulation of the AON. (H) Right: Traces showing the position of the right (green) and left (purple) eyes in response to a four pulses 10 Hz stimulation of the AON in a 161 mm larva. Data are shown as mean ± s.d. Left: Frames showing the position of the eyes (top, left eye; bottom, right eye) before (left) and after (right) stimulation in the AON. (I) Graph showing the correlation between the right and left eye movements of the 161 mm larva shown in H. Abbreviations: nIV Trochlear Motor Nucleus. px Pixels. Scale bar = 300 µm in Bi-iii.

To investigate the presence of eye movements, we first used video tracking (N = 31) analyzing the videos with DeepLabCut, an open-source software that employs artificial neural networks to estimate poses (Mathis et al., 2018). A video camera was placed facing one of the eyes of the preparation, and either the AON or the trochlear nucleus (nIV) were electrically stimulated (Fig. 1A; 4 pulses, 10 Hz). The first question was whether larvae have eye movements, and we thus started with large larvae close to metamorphosis (> 140 mm, N = 6). Reliable eye movements were evoked as can be seen in the representative example of a 145 mm larva shown in Figure 1D (Movie 1). We then investigated smaller larvae and were able to observe eye movements in larvae as small as 67 mm (Fig. 1C; N = 4). The range of movement varied depending on the individual’s development, with larger larvae exhibiting larger eye movement amplitudes than smaller ones (Fig. 1C-D). Interestingly, in larvae, a single slow movement was observed after applying four stimulation pulses (Fig. 1C-D; Movie 1), whereas late metamorphic (N = 4), postmetamorphic (N = 2), and adult lampreys (N = 1) exhibit four rapid movements, one for each stimulation pulse applied (Fig. 1E; Movie 2). These differences in eye movement velocity could be due to a low degree of development of the oculomotor circuits and/or the extraocular muscles, or rather to a mechanical constrain of the eye given the underdevelopment of the orbit. To investigate this, electromyograms (EMGs) in the dorsal rectus (DR) muscle of larvae (N = 2) were performed in response to electrical stimulation of the AON or the nIV. In Figure S1 (top trace), the EMG activity in the DR of an adult lamprey in response to a four pulses electric stimulation of the AON (10 Hz) is shown, and activity can be seen in response to each of the pulses. Below, the activity in the DR of a 161 mm larva is shown. In the case of the experiments performed in larvae, the signal recorded was weaker due to the small size of the extraocular muscle, but a clear response was observed after each of the four stimulation pulses applied in the AON. This indicates that the muscle contracts in a similar fashion to adults for each of the pulses, and that the differences in eye movement kinematics are due to the underdevelopment of the eye/orbit.

We next investigated whether eye movements in lamprey larvae occur in a coordinated manner, as observed in adults (Rovainen, 1976; Wibble et al., 2022), or if there are movements preceding this coordination. For this, we placed a video camera on top of the animal to track both eyes in response to electrical stimulation of the AON. We observed that coordinated movements after AON stimulation occur as soon as the eyes move (Movie 3). In Figure 1F, the trajectories of the eyes in response to stimulation of the AON (4 pulses, 10 Hz) are shown in a 68 mm larva, which are correlated indicating their coordinated nature (Fig. 1G; Pearson’s correlation analysis; r = –0.784; p < 0.001; Movie 3). This coordination was observed in all recorded larvae larger than 67 mm (N = 23), although this correlation was higher in large pre-metamorphic larvae (Fig. 1H-I; Pearson’s correlation analysis; r = 0.953; p < 0.001; Movie 4). Altogether, these results demonstrate that, despite living borrowed and having underdeveloped eyes covered with skin, lamprey larvae have coordinated eye movements as soon as their extraocular muscles (and their innervation; see Pombal et al., 1994) develop.

### VOR is the first type of eye movement that emerges

The VOR probably emerged as the first reflex for gaze stabilization, would therefore be evolutionarily the earliest, and the first to appear in development (Straka and Baker, 2013). In lamprey larvae, vestibular information from the AON is initially directed to the SC when the larvae are 27-30 mm long. As they grow larger (> 60 mm), the AON axons project ventrorostrally and eventually form synapses with the nIII, located in the ventral midbrain (Pombal et al., 1994, 1996; Fritzsch, 1998). Interestingly, this was the smallest size at which we could detect eye movements, and no eye movements could be evoked stimulating the nIII, nIV or nVI nerves in smaller animals that did not respond to AON stimulation. This suggests that the VOR is the first type of eye movement that appears in lampreys. On this basis, we decided to analyze the VOR nature of the first eye movements in ⁓60 mm larvae. For this, we first confirmed that eye movements elicited by AON stimulation are the result of activating the vestibular system by monitoring eye movements in response to mechanical stimulation of the labyrinth in larvae > 60 mm (Fig. 2A; N = 3). This resulted in eye movements (Fig. 2B-C; Movie 5), like those evoked by electrical stimulation of the AON (see above), confirming their VOR nature.

**Fig. 2.**
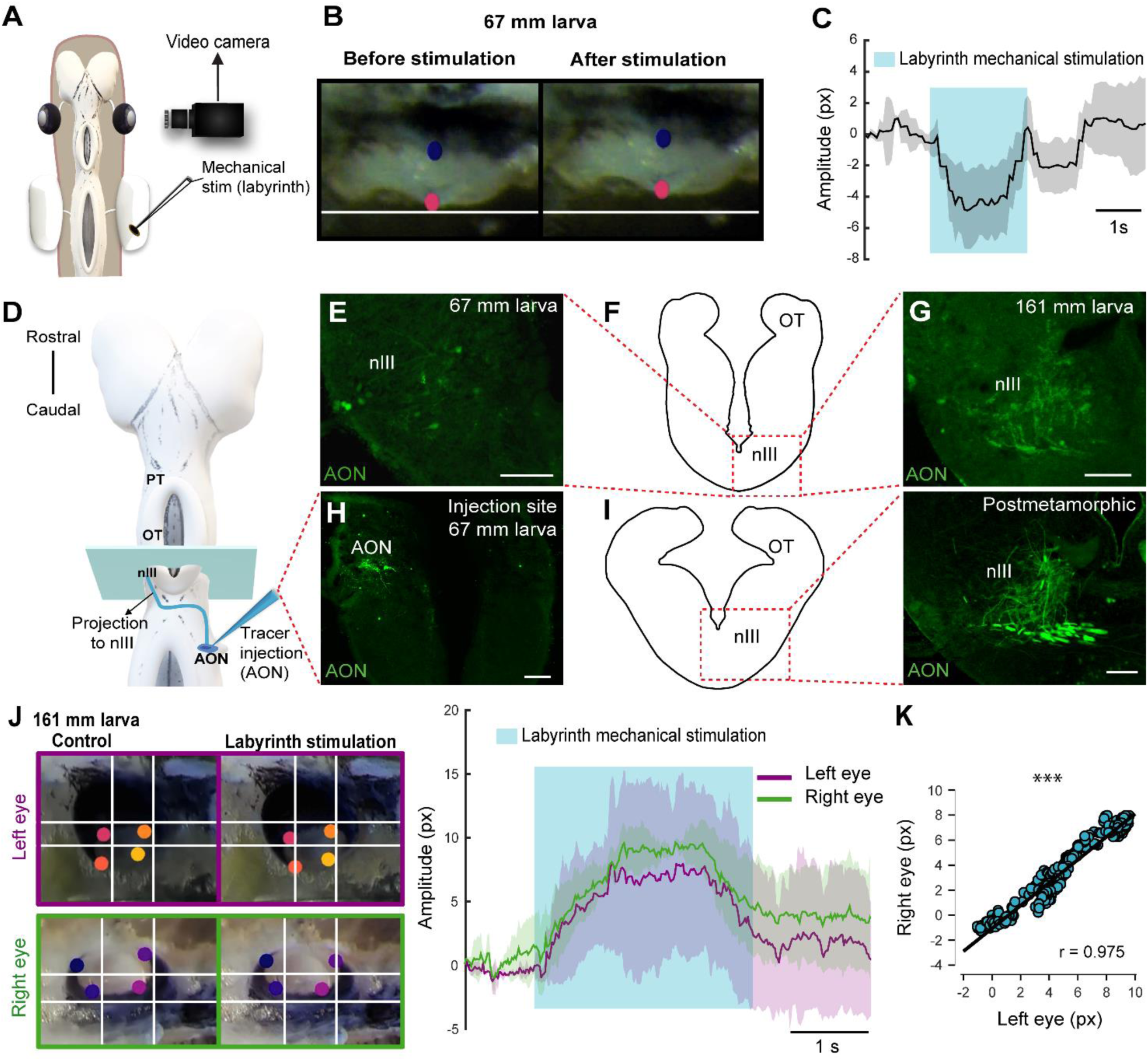
Lamprey larvae exhibit VOR. (A) Schematic showing the *ex vivo* preparation used to monitor eye movements in response to mechanical stimulation of the labyrinth. (B) Eye of a 67 mm larva before (left) and after (right) vestibular stimulation. Colored dots are the labels used to track eye movements. (C) Graph showing the eye movement of the same larva after labyrinth stimulation. The blue shaded area indicates the duration of the vestibular stimulation. (D) Schematic dorsal view of the lamprey larva brain showing the location of the tracer injection site (Anterior Octavomotor Nucleus, AON) and its anterograde projections (blue line) reaching the contralateral oculomotor nucleus (nIII). (E-I) The level of the drawings shown in F and I (left) corresponds to the blue rectangle in D and the dotted rectangle in both drawings indicates the location of the photomicrographs at the level of the contralateral nIII showing axons anterogradely labeled from the AON in a 67 (E) and a 161 (G) mm larva and a posmetamorphic animal (I right). (H) Representative injection site in the AON of a 67 mm larva. (J) Left: Representative frames showing eye position (top, left eye; bottom, right eye) before (left) and after (right) vestibular stimulation. Right: Traces showing the position of the right (green) and left (purple) eyes in response mechanical stimulation of the labyrinth in a 161 mm larva. The blue shaded area denotes the duration of the vestibular stimulation. (K) Graph showing the correlation between the movements of the right and left eyes. In all graphs, data are shown as mean ± s.d. Abbreviations: OT Optic Tectum, PT Pretectum, px Pixels. Scale bar = 100 µm in E and H; 50 µm in G and I.

To test if the appearance of VOR becomes functional as soon as the underlying neural circuits appear or if it emerges later, tracer injections were performed in the AON (N = 12) to determine the larval size at which the VOR circuit develops (Fig. 2D-I). Tracer injections into the AON confirmed the projections to the nIII previously shown both in adults and larvae lampreys (Pombal et al., 1996; Fritzsch, 1998; Wibble et al., 2022). Anterogradely labeled fibers were found in the nIII of > 60 mm larvae (Fig. 2E-H; N = 6) that also showed eye movements in response to AON stimulation, but no anterogradely labeled fibers were found in smaller larvae (< 60 mm, N = 4; not shown). The AON projection to the nIII increased considerably in larger larvae, postmetamorphic (N = 1) and adult lampreys (N = 1; Fig. 2F,G,I). The increase in fibers number throughout development was also observed at their crossing point, caudal to the nIII (Fig. S2A-C). Next, we tested whether the coordinated eye movements observed in response to electric stimulation of the AON could also be evoked in response to mechanical stimulation of the labyrinth, confirming its VOR nature (N = 2). In Figure 2J-K a representative example is illustrated of a 161 mm larva that shows a significant correlation between the movement of the right and left eyes (Pearson’s correlation analysis; r = 0.975; p < 0.001), like those observed in the same stage of large pre-metamorphic larvae after electric stimulation of the AON (see above). These results show that VOR is the first type of eye movements that emerges.

### Lamprey larvae exhibit eye movements in response to light stimulation

The presence of VOR in larval lampreys raised the question of whether other types of eye movements also appear during this period. In lampreys, visual information can generate eye movements mostly through three visual centers: the PT that mediates the optokinetic reflex and has been also suggested to mediate phototactic responses, the OT that is thought to mediate goal-oriented movements, and pallial regions thought to be the precursor of cortical regions (see above). Thus, we first performed electric stimulation of these areas to check whether eye movements could be evoked (N = 26). Eye movements were observed in larvae as small as 100 mm in response to both pretectal (N = 7; Fig. 3A) and tectal stimulation (N = 5; Fig. 3B). Interestingly, no eye movements could be evoked in these larvae stimulating the visual area in pallium, nor activity could be evoked in the middle rhombencephalic reticulospinal nucleus (MRRN), and the same happened in metamorphic and recent postmetamorphic animals, although activity in the MRRN and/or eye movements were observed after tectal/pretectal stimulation (see below), in agreement with the late development of tectal and pretectal motor outputs (see below).

**Fig. 3.**
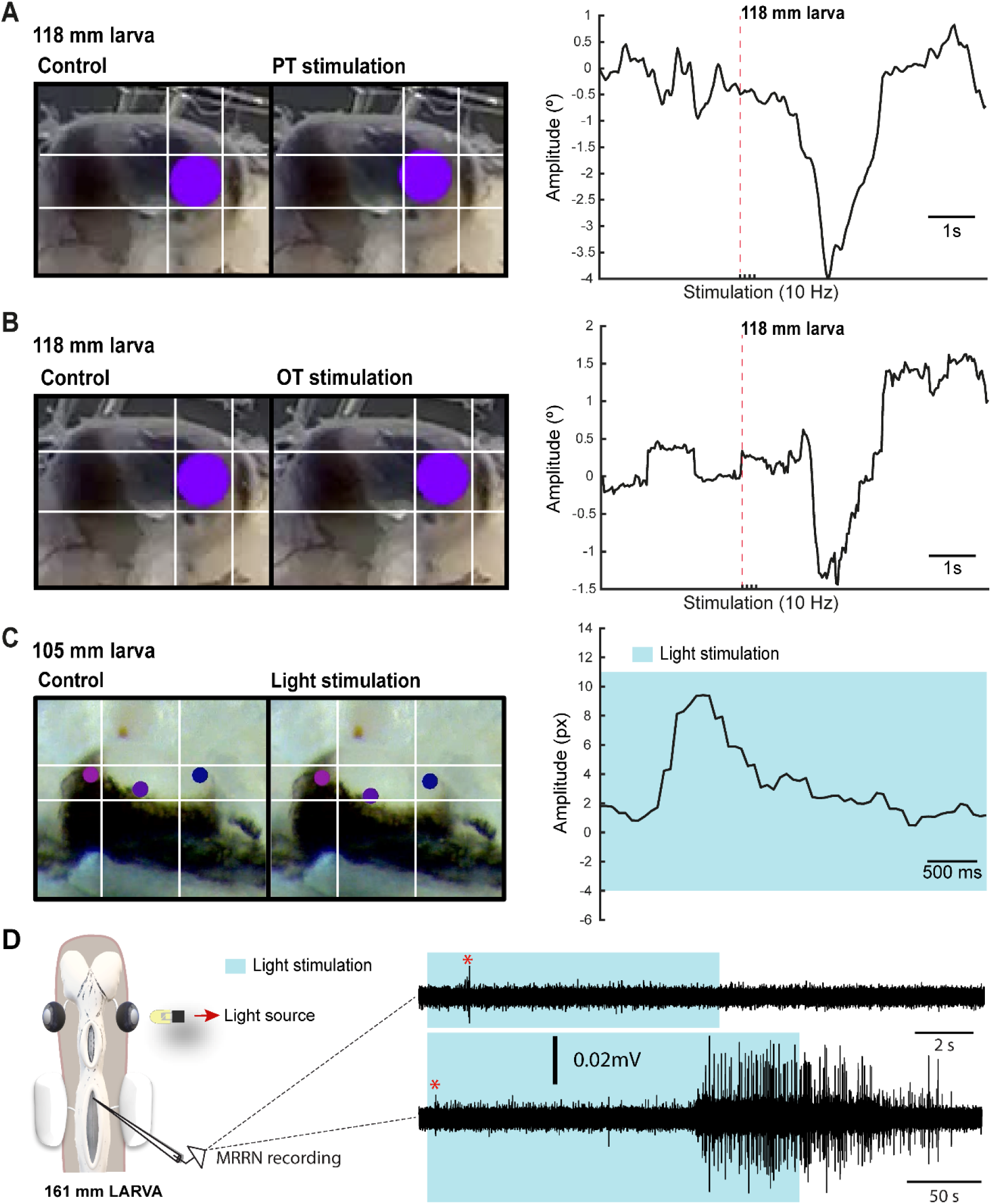
Light-evoked responses. (A) Left: Eye of a 118 mm larva before (left) and after (right) a four pulses electric stimulation of the pretectum (PT; 10 Hz). Right: Graph showing the eye movement in response to the stimulation. In (B) the eye movement is shown in response to stimulation of the optic tectum (OT; 10 Hz). The red dotted line in A and B indicates when the first pulse was applied. (C) Representative frames (left) showing the eye position before and after light stimulation for the eye movement shown in the graph (right). The shadowed blue area indicates light stimulation to both eyes. Colored dots indicate the labels used for tracking eye position. (D) Representative traces showing extracellular activity in the middle rhombencephalic reticulospinal nucleus (MRRN) of a 161 mm larva in response to a 10 s (top trace) and to a 50 s (bottom trace) light stimulation presented to one eye. The experimental setup is shown in the schematic to the left. The shadowed blue area denotes the duration of the stimulation, whereas the red asterisk signals the short latency evoked responses. Abbreviations: px Pixels.

The presence of eye movements evoked from retinorecipient areas after electric stimulation raised the question of whether visual stimuli could also trigger eye movements. The low degree of development of the retina, with only the central retina differentiated, indicates that only basic visual features can be integrated at larval stages (Suzuki and Grillner, 2018), and we therefore analyzed eye movements in response to light stimuli (N = 11). Although no eye movements were observed with light stimuli presented with a LED to one eye, these were evoked in response to broad light stimulation to both eyes after a period of darkness in larvae larger than 100 mm (Fig. 3C; N = 5). Given the presence of light evoked eye movements, we also tested whether extracellular activity could be evoked in the MRRN, indicative of phototactic responses (see below). Weak short latency activity was observed in larvae larger than ⁓100 mm in response to light stimulation presented with a LED to one eye (Fig. 3D, red asterisk; N = 2). Furthermore, a strong activation was evoked when light stimulation was maintained several seconds (Fig. 3D, bottom trace). These results indicate that visual information can generate eye and body movements already in larvae, although their nature and functional relevance remains to be determined, and likely basic phototactic responses mediated by the eyes. The presence of light-evoked eye movements suggested that OKR might be also present in larvae. To investigate this, we did tracer injections in the PT (N = 28) to see whether anterogradely labeled fibers were present in the nIII. However, no evidence of a PT to nIII connection was found (not shown), indicating that the anatomical substrate of OKR (Wibble et al., 2022) is missing at this stage.

### Development of tectal and pretectal projections to the MRRN in parasitic lampreys

In adult lampreys, neurons in the deep layer of the OT send direct projections to the brainstem, making synaptic contacts onto reticulospinal neurons in the MRRN. Tectal projections are both contra– and ipsilateral and control orienting and evasive responses, respectively (Kardamakis et al., 2015; 2016; 2017, Suzuki et al., 2019). The PT also sends direct projections to the reticulospinal neurons in the MRRN as well as to the posterior rhombencephalic reticulospinal nucleus, providing an additional substrate for the visually evoked motor responses in lampreys (Zompa and Dubuc, 1996; 1998; Ullén et al., 1997; Capantini et al., 2017). However, when these circuits appear during development is not yet known. Thus, we combined extracellular recordings in the MRRN in response to stimulation of the OT and the PT at different developmental stages with tracer injections in this region to uncover when these connections are established. Given that the stepwise development of the visual system observed in lampreys is consequence of their very long larval period, we speculated that differences in the visual circuits controlling orienting and evasive responses might be present in lampreys with different lifestyles. Thus, we compared the tectal/pretectal inputs of a parasitic lamprey (*Petromyzon marinus*), with those of the non-parasitic Northern Japanese brook lamprey (*Lethenteron* sp).

In parasitic lampreys, no evidence of projections from either OT or PT to the MRRN was found in larvae. Neurobiotin injections in the MRRN were performed (N = 12) and in all cases the tectal and pretectal regions were devoid of retrogradely labeled neurons (Fig. 4A,C). In agreement with this, no activity was evoked in the MRRN after stimulation of the PT (Fig. 4B, red trace). Interestingly, in a couple of larvae short latency responses were evoked in the MRRN in response to high intensity stimulation pulses in the PT (not shown) despite the lack of retrogradely labeled neurons from the MRRN, most likely due to the activation of the nMLF dendrites shown to reach this area (Suzuki et al., 2015a, b). On the other hand, responses were evoked in the MRRN of larvae larger than 60 mm after tectal stimulation although only with long latency onsets, indicative of a polysynaptic pathway (Fig. 4B, D, green trace, and E: response onset = 30.68 ± 1.29 ms; n = 5). These responses were initially potentiating, but usually decayed after the third pulse, as can be seen in the size of the evoked responses for a four pulses 10 Hz stimulation (Fig. 4F).

**Fig. 4.**
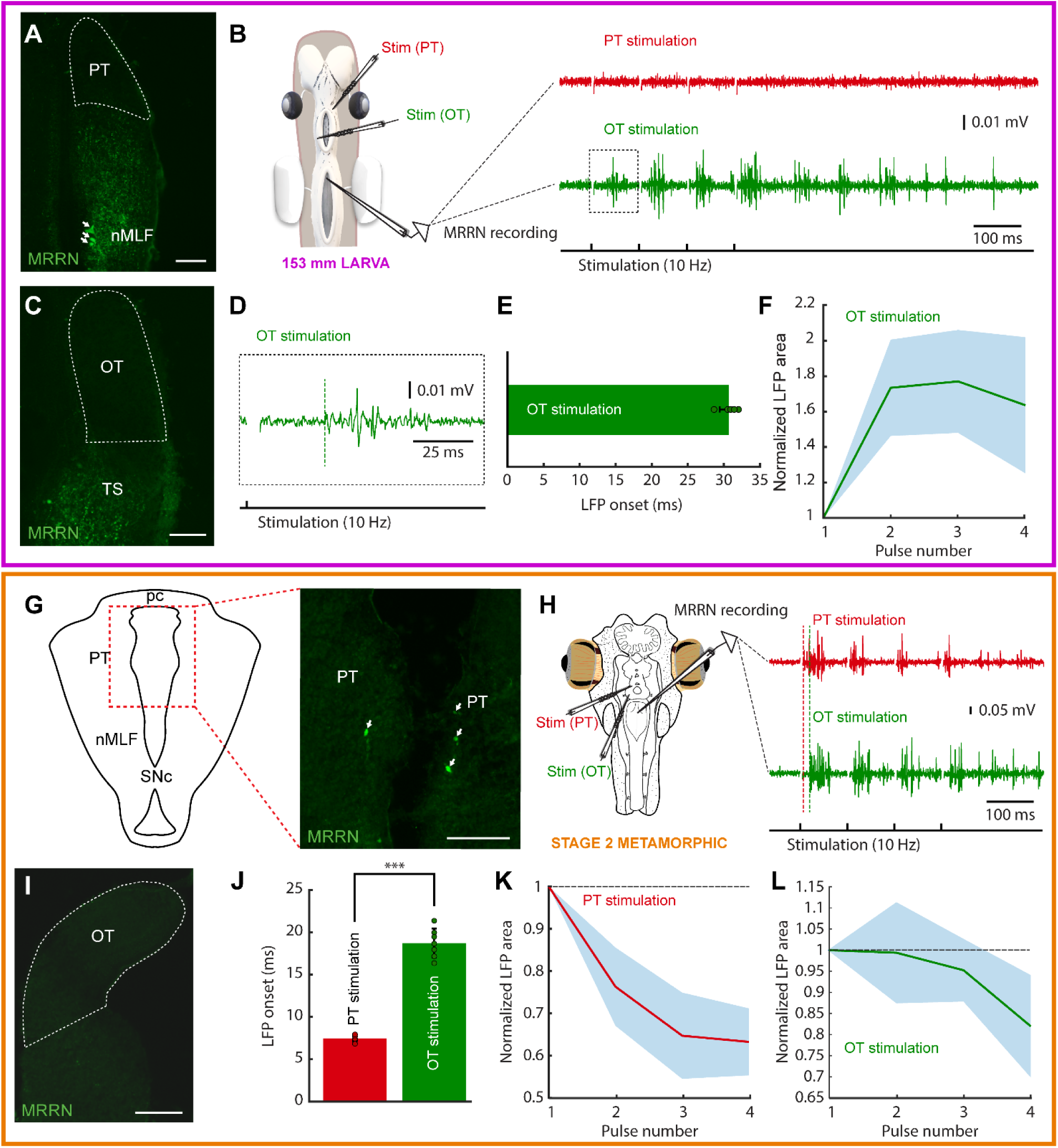
Tectal and pretectal motor outputs in larvae/early metamorphic. The colored rectangles group results belonging to the developmental stage indicated under the schematic of the experimental preparation (magenta: 153 mm larva; orange: stage 2 metamorphic). (A) Photomicrograph showing the lack of retrogradely labeled neurons in the pretectum (PT, indicated by a dashed line) after a dextran injection in the middle rhombencephalic reticulospinal nucleus (MRRN) of a 153 mm larva. Retrogradely labeled neurons can be seen in the nucleus of the medial longitudinal fasciculus (nMLF; arrows). (B) Schematic showing the preparation used to record neuronal activity in the MRRN of the same larva (left). Electric stimulation of the PT did not evoke activity in the MRRN (right, red trace), whereas electric stimulation of the optic tectum (OT) resulted in extracellular activity in response to each of the four applied pulses (right, green trace). (C) No retrogradely labeled neurons were observed in the OT after a dextran injection in the MRRN (surrounded by a dashed line). (D) Magnified view of the region indicated by a dashed line square in the trace shown in B indicating the onset of the evoked response (vertical dashed green line). (E) Plot showing the average onset times of the responses evoked in the MRRN after electric stimulation of the OT. (F) Mean responses in the MRRN in response to 4 pulses electric stimulation of the OT (10 Hz), combining data of 5 larvae ranging from 65 to 161 mm. (G) Schematic drawing (left) indicating the location of the photomicrograph (right) showing bilateral retrogradely labeled MRRN projecting neurons in the periventricular aspect of the PT in a stage 2 metamorphic lamprey (arrows). (H) Left, schematic showing the preparation used to record extracellular activity in the MRRN (right) in response to electric stimulation of the PT (red trace), and the OT (green trace). Vertical dashed lines indicate the onset of the evoked responses. (I) Photomicrograph showing the lack of retrogradely labeled neurons in the OT after a tracer injection in the MRRN. (J) Graph showing the difference between response onsets in the MRRN after PT (red) and OT (green) stimulation. (K-L) Graphs showing the mean MRRN activity in a stage 2 animal evoked by stimulation of the PT (K) and OT (L). Values are normalized to the first local field potential (LFP). In the electrophysiological traces, stimulation artifacts were removed for clarity. In all graphs, data are shown as mean ± s.d. Abbreviations: pc Posterior commissure, SNc Substantia Nigra pars compacta, TS Torus Semicircularis. Scale bar = 100 µm in A, C, G and I.

Given the lack of tectal and pretectal projections to the MRRN in larvae of parasitic lampreys, we next analyzed their presence in metamorphic animals (N = 7). Tracer injections in the MRRN resulted in retrogradely labeled neurons in the PT already in the earliest metamorphic stage that we could analyze (stage two, Youson and Potter, 1979; Fig. S3A-C). Although the pretectal projection to the MRRN was developed, only some retrogradely neurons were found in the periventricular region (Fig. 4G). Still, clear activity could be recorded in the MRRN after electric stimulation of the PT (Fig. 4H, red trace; 10 Hz, four pulses). Noticeably, anterogradely fibers were observed at this and later stages (N = 6) in the nIII after tracer injections in the PT, showing that the OKR substrate also develops early during metamorphosis (not shown). However, no retrogradely labeled neurons were found in the OT after tracer injection in the MRRN (Fig. 4I). In agreement with the anatomical experiments, the electrophysiological activity recorded in the MRRN after pretectal stimulation showed a short latency (7.43 ± 0.37 ms), significantly shorter than that evoked after tectal stimulation (18.64 ± 1.71 ms; Fig. 4J; unpaired *t* test, p < 0.0001; n = 14), thus confirming the direct pretectal projections to the MRRN and the lack of monosynaptic tectal projections.

In later metamorphic stages (stage five and later), tracer injections in the MRRN resulted in retrogradely labeled neurons both in periventricular and lateral portions of the PT, in a situation similar to that reported in adults (Fig. 5A; Zompa and Dubuc, 1996; El Manira et al., 1997; Capantini et al., 2017). The conspicuous population of fusiform neurons extending their dendrites into the optic tract reported in adults could also be observed at this stage (Fig. 5A, arrows). Extracellular recordings in the MRRN in response to pretectal stimulation resulted in short latency responses (6.55 ± 0.46 ms; Fig. 5B, red trace), in agreement with a monosynaptic input (von Twickel et al., 2019). Contrary to PT, no retrogradely labeled neurons were found in the OT after tracer injection in the MRRN (Fig. 5C, dashed area). Accordingly, although extracellular activity could be consistently evoked in the MRRN in response to tectal stimulation (Fig. 5B, green trace), these responses had a long latency (22.5 ± 1.94 ms), significantly longer than the pretectal evoked responses (unpaired *t* test, p < 0.0001; Fig. 5B, see vertical dashed lines, and 5D; n = 12), indicative of a polysynaptic pathway and similar to the situation found in larval stages (see above). Interestingly, only in late metamorphic/early postmetamorphic lampreys retrogradely labeled neurons could be detected in the OT, and still in small numbers (Fig. 5G). Accordingly, weak responses with short latencies were observed in the MRRN after tectal stimulation, followed by a big response component, which corresponds to the polysynaptic response observed in previous stages (Fig. 5H). Contrary to larvae, in metamorphic animals the MRRN responses after tectal or pretectal stimulation were not consistently potentiating, and both inhibitory (Fig. 4K, L) and potentiating cases (Fig. 5E, F) were found after a four pulses stimulation (10 Hz). The presence of polysynaptic connections in larvae suggests that visual stimuli integrated by the OT and the PT can evoke motor responses through the reticular formation, most likely related to phototactic responses, as suggested by the above presented experiments using light stimulation. However, in parasitic lampreys, the full map of projections is not completely developed until very late stages of the metamorphosis, with the beginning of their parasitic and migratory feeding life.

**Fig. 5.**
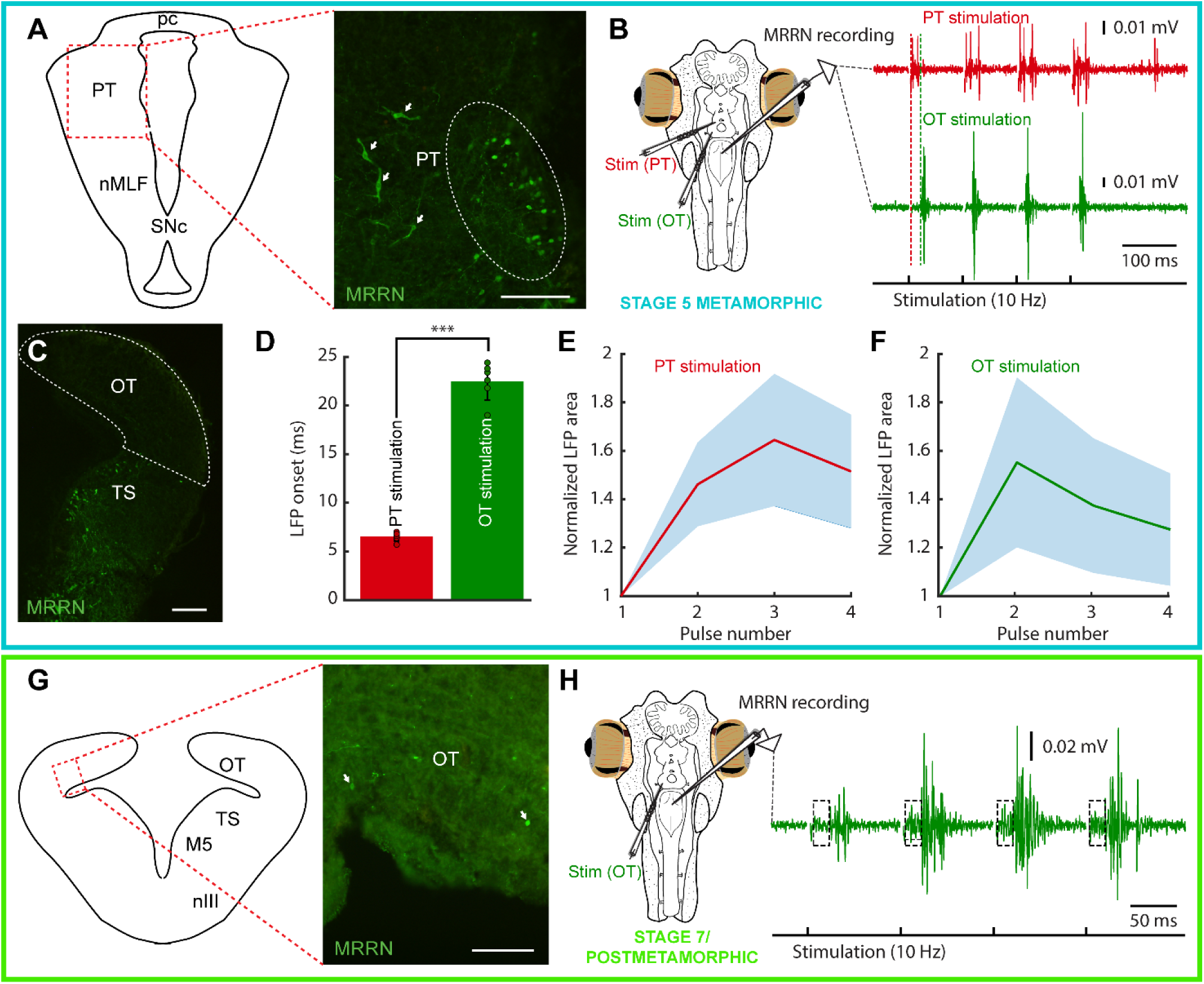
Tectal and pretectal projections to the MRRN in metamorphic and postmetamorphic lampreys. The colored rectangles group results belonging to the developmental stage indicated under the schematic of the experimental preparation (blue: stage 5 metamorphic; green: stage 7 metamorphic/postmetamorphic). (A) Schematic drawing (left) indicating the location of the photomicrograph (right) showing retrogradely labeled neurons in the pretectum (PT) of a stage 5 metamorphic lamprey after a Neurobiotin injection in the middle rhombencephalic reticulospinal nucleus (MRRN). Projection neurons can be observed both in the periventricular region (dashed line oval), and in lateral aspects (arrows). (B) Extracellular responses in the MRRN after stimulation of the PT (red trace) and optic tectum (OT, green trace). The onset of the extracellular activity is indicated by a dashed red line for PT stimulation, and a dashed green line for OT stimulation. (C) Photomicrograph showing that no retrogradely labeled neurons can be seen in the OT (indicated by a dashed line) of a stage 5 metamorphic lamprey after a Neurobiotin injection in the MRRN. (D) Graph showing that the onsets of MRRN responses evoked by PT stimulation (red) were significantly shorter than those evoked by OT stimulation (green; unpaired t-test). (E-F) Graphs showing the mean responses evoked in the MRRN of a stage 5 metamorphic animal evoked by stimulation of the PT (E) and OT (F) in response to 4 pulses (10 Hz). Values are normalized to the first local field potential (LFP). (G) Schematic drawing (left) indicating the location of the photomicrograph (right) showing a few retrogradely labeled neurons from the MRRN (arrows) in the OT of a late metamorphic animal (stage 7/recent postmetamorphic). (H) Extracellular responses in the MRRN of a late metamorphic animal (stage 7/recent postmetamorphic) in response to OT stimulation (4 pulses, 10 Hz). A two-components response can be observed: a fast onset weak response (indicated by a dashed line rectangle) followed by a stronger signal. In the electrophysiological traces, stimulation artifacts were removed for clarity. In all graphs, data are shown as mean ± s.d. Abbreviations: nMLF Nucleus of the Medial Longitudinal Fasciculus, pc Posterior commissure, SNc Substantia Nigra pars compacta, TS Torus Semicircularis, nIII Oculomotor Nucleus, M5 Retinopetal Nucleus of Schöber. Scale bar = 100 µm in A, G and M; 200 µm in C and I.

### Earlier development of tectal projections in non-parasitic lampreys

The late appearance of the tectal and pretectal motor projections in parasitic lampreys indicates that the development of the visual system is completed in parallel with the biological needs of the animal. This raised the question of whether tectal and pretectal motor outputs develop differently in non-parasitic lampreys. Thus, we investigated tectal and pretectal projections in a non-parasitic, landlocked species (Northern Japanese brook lamprey, *Lethenteron* sp. N). Whole-brain confocal imaging allowed us to confirm that the optic nerve fibers did not project to the OT but predominantly project to the PT in larvae smaller than 70 mm (Fig. 6A-B), as previously reported (de Miguel et al., 1990; Cornide-Petronio et al., 2011). Nevertheless, retrograde labeling in these small larvae already showed that there were tectal as well as pretectal neurons projecting to the ipsilateral MRRN (N = 3/3; Fig. 6A-C). In larvae larger than 70 mm in body length, contralaterally-projecting tectal neurons were also found at least in some specimens (N = 2/5; Fig. 6D-H). At the same time, tracer injections into the OT (Fig. 6I) resulted in anterogradely labeled fibers of tectal neurons that projected to the MRRN region in both large larvae (Fig. 6J; N = 3) and postmetamorphic animals (not shown; N = 4). The labeled fibers in larvae were in close contact with the MRRN Müller neurons (Fig. 6K), indicating that the tectal neurons are connected directly to the MRRN. Lastly, in the postmetamorphic animals, both ipsi– and contralaterally projecting tectal neurons were observed (N = 4/4; Fig. 6L-N,P). These labeled neurons were all in the deep layer (DL) of the OT (Fig. 6C,G,H,O). Additionally, pretectal neurons projecting to the MRRN were also labeled on both ipsi– and contralateral sides in postmetamorphic animals (N = 4/4; Fig. 6Q). These results show that the differentiation of the tectal projections to the MRRN is accelerated in the non-parasitic, landlocked lampreys compared to parasitic species.

**Fig. 6.**
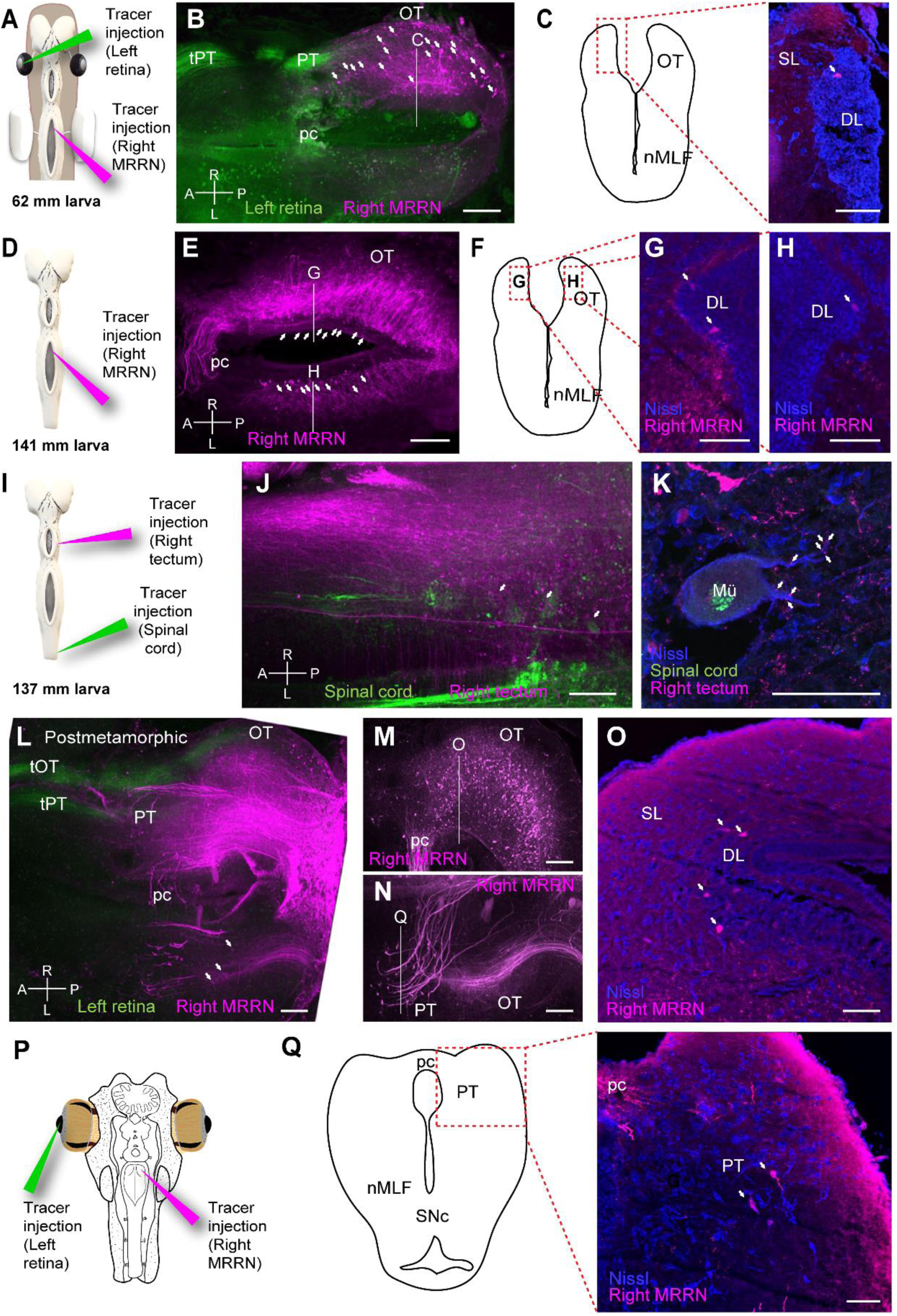
Tectal and pretectal projections to the MRRN in non-parasitic lampreys. (A) Schematic drawing indicating the location of the tracer injections sites in the left retina and the right middle rhombencephalic reticulospinal nucleus (MRRN). (B) Whole-brain confocal image showing dual labeling of the optic nerve fibers (green) and MRRN projecting tectal neurons (magenta) in a 62 mm larva, dorsal view. The optic nerve predominantly projects to the pretectum (PT) via the pretectal tract (tPT). The PT is located at the same level of the posterior commissure (pc) on the anterior-posterior axis. The optic tectum (OT) contains retrogradely labeled neurons in its rostrocaudal extent (arrows). (C) Schematic drawing (left) indicating the location of the photomicrograph (right) at the level depicted in B showing a retrogradely labeled neuron in the OT of a 62 mm larva. The labeled neuron is not in the superficial (SL) but in the deep periventricular layer (DL, arrow). (D) Schematic drawing indicating the location of the tracer injection site in the MRRN. (E) Whole-brain confocal image showing MRRN-projecting tectal neurons (magenta) in a 141 mm larva, dorsal view. Labeled neurons are observed on both sides of the OT (arrows). (F-H) Schematic drawing and photomicrographs at the levels depicted in E showing retrogradely labeled neurons in the right (G) and left (H) OT in a 141 mm larva. The labeled neurons are present in the DL (arrows). The location of the photomicrographs in G and H is shown in the schematic drawing in F. (I) Schematic drawing indicating the location of the tracer injection sites in the right OT and the spinal cord (SC). (J) Whole-brain confocal image showing dual labeling of retrogradely labeled SC-projecting cells (green) and anterogradely labeled tectal fibers (magenta) in a 137 mm larva, dorsal view. Numerous MRRN-projecting tectal fibers are observed (arrows). (K) Photomicrograph showing anterogradely labeled tectal terminals on a large Müller neuron (Mü) in the MRRN region, suggesting direct connections between them. (L) Whole-brain confocal image showing dual labeling of the optic nerve fibers (green) and MRRN-projecting tectal neurons (magenta) in postmetamorphic animals, dorsal view. In addition to tPT, the tectal tract (tOT) of the optic nerve is observed. Some contralateral MRRN-projecting tectal neurons are also distinguished (arrows). (M, N) Region-specific images for the right (M) and left (N) OT, generated by different Z-stack projections to visualize ipsilateral MRRN-projecting tectal neurons and contralateral MRRN-projecting pretectal neurons, respectively. (O) Photomicrograph at the level depicted in M showing retrogradely labeled neurons from MRRN in the DL (arrows) of the ipsilateral OT. (P) Schematic drawing indicating the location of the tracer injection sites in the left retina and the right MRRN. (Q) Schematic drawing (left) indicating the location of the photomicrograph (right) at the level depicted in N, showing retrogradely labeled neurons from MRRN in the contralateral PT (arrows) in postmetamorphic animals. Abbreviations: A Anterior, L left, nMLF Nucleus of the Medial Longitudinal Fasciculus, P posterior, R right, SNc Substantia Nigra pars compacta. Scale bars = 100 µm in B, E, J, L, M, and N; 50 µm in C, G, H, K, O, and Q.

### The nMLF mediates the first tectal-evoked motor responses

Although our results show that direct tectal and pretectal projections to the MRRN are not present in larvae of parasitic lampreys, the OT can evoke activity in the MRRN via polysynaptic pathways (see above). The main candidate to mediate these motor outputs from the OT is the nMLF. This region sends motor commands to the brainstem and the SC and has been suggested to generate negative phototactic responses (Suzuki and Grillner, 2018). Tracer injections in the MRRN of larvae and early postmetamorphic lampreys showed that the nMLF is the only visual region that projects to the MRRN (Fig. 7A-B; N = 16; see also Fig. 4A). The other populations projecting to the brainstem were found in ventral aspects of the thalamus, most likely belonging to the diencephalic locomotor region (El Manira et al., 1997; not shown), and the mesencephalic locomotor region (Grillner and El Manira, 2020; not shown). Thus, we tested whether the nMLF mediates the polysynaptic responses evoked from the OT. For this, we performed electric stimulation of this region in metamorphic animals (Fig. 7C; 10 Hz, four pulses) and recorded the extracellular activity in the MRRN before (black trace) and after lesioning the nMLF (red trace; N = 2). The long latency evoked responses drastically reduced after the lesion (Fig. 7C-D; unpaired *t* test, p < 0.0001; n = 30), indicating that the nMLF is the main structure that mediates the downstream motor responses of the OT.

**Fig. 7.**
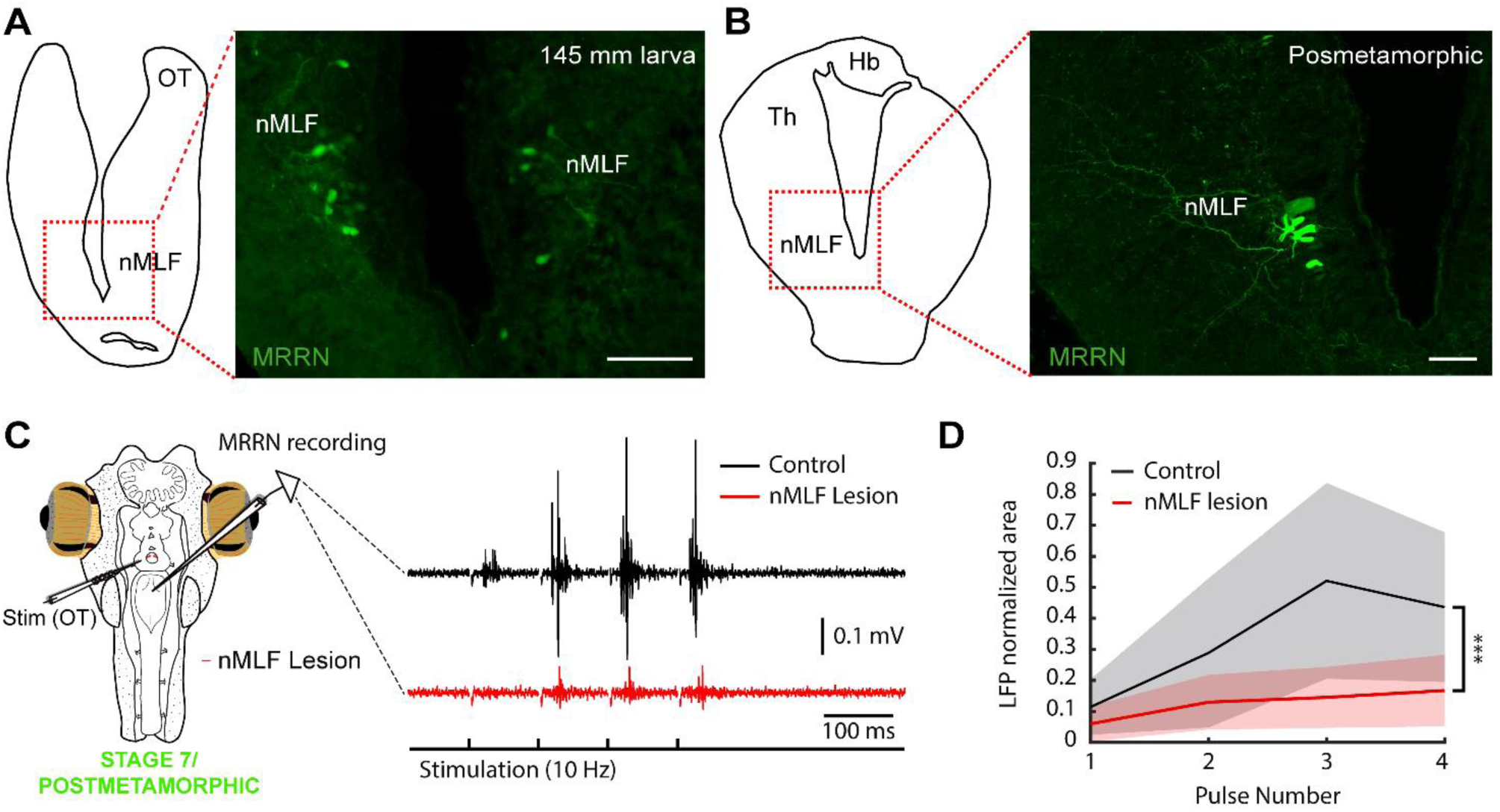
The nMLF mediates tectal responses in larvae and metamorphic lampreys. (A) Schematic drawing (left) indicating the location of the photomicrograph (right) showing bilateral retrogradely labeled neurons in the nucleus of the medial longitudinal fasciculus (nMLF) of a representative 145 mm larva after a tracer injection in the middle rhombencephalic reticulospinal nucleus (MRRN). (B) Schematic drawing (left) indicating the location of the photomicrograph (right) showing MRRN projecting neurons in the nMLF of a postmetamorphic animal. (C) The polysynaptic responses evoked in the MRRN after electrical stimulation of the optic tectum (OT; black trace) are drastically reduced after lesioning the nMLF in a stage 7/recent postmetamorphic (red trace). Stimulation artifacts were removed for clarity. (D) Graph showing the significant reduction in MRRN activity in response to OT stimulation after lesioning the nMLF. Abbreviations: Hb Habenula, Th Dorsal Thalamus. Scale bars = 100 µm in A and B.

## Discussion

Lampreys possess a well-developed visual system, with fundamental characteristics found in all vertebrates, whose components and retinofugal pathways develop in a stepwise manner during a long larval period (Suzuki and Griller, 2018). Here we show that the visuomotor pathways also develop stepwise and that, interestingly, the oculomotor system in lamprey larvae becomes functional long before the development of its visual function, as we show that eye movements are present already in larvae. Previous studies have shown that in first instance the vestibular information is directed to the SC to generate body motor behaviors and, when lamprey larvae are larger, AON axons project to the nIII (Pombal et al., 1994; Pombal et al., 1996). These studies and our tracer injections show that the appearance of VOR corresponds temporally with the establishment of its underlying neural circuits. Interestingly, some degree of eye coordination also appears as soon as VOR emerges, while completely synchronized eye movements are only observed in large larvae close to metamorphosis. Larval eye movements exhibit slow kinematics compared to the adult ones. However, EMG activity in the extraocular muscles shows that their responses are similar to adults, meaning that the improved eye movement performance is consequence of the eye/orbit development. The number of projections is more evident in larger larvae, which accordingly display a larger range of motion than small larvae, although larger eye movement amplitudes are likely dependent on an improved muscle function too (Easter and Nicola, 1997). Interestingly, not only VOR but also light evoked eye movements are present in > 100 mm larvae. This could be related with the minimum larval size needed to enter metamorphosis, as previously suggested (Pombal et al., 1994).

It is curious to imagine the utility of such an early development of eye movements. Lamprey larvae live buried in the mud on which they feed (Hardisty, 2006) so they do not hunt, and their eyes are covered by skin and are not functional image-forming structures (Kleerekoper, 1972; Suzuki and Grillner, 2018). Regarding VOR, it is hard to imagine any behavioral relevance. However, given that both electric stimulation of the PT/OT and light stimulation evoke eye movements, it is possible that light-evoked ocular movements play a role in phototactic responses. It has been shown that retinal projections to the PT directly target nMLF neurons, suggesting that this pathway can generate negative phototaxis in larvae (Suzuki et al., 2015a, b). Interestingly, our results show a consistent activation of the MRRN via tectal stimulation, which is mediated via the nMLF. Given the MRRN activity evoked by light in our experiments, it is likely that the OT can also generate phototactic responses through the nMLF. The presence of a basic retinotopy already in the larval OT (Cornide-Petronio et al., 2011) could provide a substrate so that motor responses can be evoked with a certain directionality, and therefore light-evoked eye movements may have a behavioral relevance, although this remains to be tested behaviorally.

It is believed that the first ocular movement that appeared during evolution is the VOR, and that goal-oriented gaze movements evolved from stabilizing responses (Walls 1962; Land, 2015). Our results show that the first eye movements during lamprey development are VOR-like followed by phototactic responses in larval stages, and then the OKR followed by goal-oriented responses emerge during metamorphosis, as indicated by the late development of pretectal and tectal motor projections. Remarkably, our results indicate that the pallial control of visuomotor responses appears even later (Fig. 8). The development of stabilizing gaze movements first, followed by goal-oriented responses is also observed in other vertebrates (Dayton et al., 1964; Horn et al., 1986; Easter and Nicola, 1997; Huang and Neuhauss, 2008; Branoner et al., 2016; Donovan et al., 2020; Forsthofer and Straka, 2023). For instance, in zebrafish the OKR and the VOR emerge shortly after hatching (73-74 h after fertilization), while spontaneous saccades appear later (between 81 and 96 h after fertilization; Easter and Nicola, 1997; Huang and Neuhauss, 2008). The development of the lamprey visual system thus follows the same pattern as in other vertebrates, albeit in a stepwise manner and during a long period spanning several years in agreement with its peculiar life cycle. Interestingly, only the VOR and basic phototactic responses emerge during the larval period whereas the tectal and pretectal motor outputs, underlying the OKR and goal-oriented responses, appear during the metamorphosis (Fig. 8). It is known that visual experience is necessary for normal development of visuomotor circuits, including the tectal motor maps (du Lac and Knudsen, 1991; Wang et al., 2015), and lamprey development also reflects this. The availability of vestibular and rough light stimuli during the larval period would allow an earlier development of VOR and basic phototactic responses. However, the motor circuits underlying OKR and visually evoked goal-oriented behaviors only develop after the eye can form images and lampreys abandon their buried lifestyle, so that the necessary sensory information is available. Interestingly, we have seen that in non-parasitic lampreys the development of these circuits is earlier than in parasitic ones, with PT to MRRN and OT to MRRN connections already appearing in larvae. In parasitic lampreys, which will perform more advanced hunting behaviors, the PT-MRRN connection appears during metamorphosis and the OT-MRRN connection is not found until the last metamorphic stage, and even at this stage the number of tectal neurons that project to the brainstem is very scarce. Thus, it seems that the OT circuits involved in goal-oriented responses are only completed after metamorphosis, when lampreys are exposed to the appropriate visual stimuli in their downstream journey in parasitic lampreys, and as soon as they start swimming in non-parasitic ones. We have not found any evidence of a pallial contribution to gaze responses even in recently transformed animals, suggesting similar sensory requirements and that its development is completed even later, most likely because of its involvement in advanced hunting behaviors. In non-parasitic lampreys, this process is accelerated likely because of their less complex use of the visual system, so that the sensory requirements for its development are also fewer. Future studies analyzing the degree of development of these circuits in postmetamorphic parasitic lampreys at different moments of their predatory phase will provide interesting insights about the sensory requirements on the development of visuomotor circuits.

**Fig. 8.**
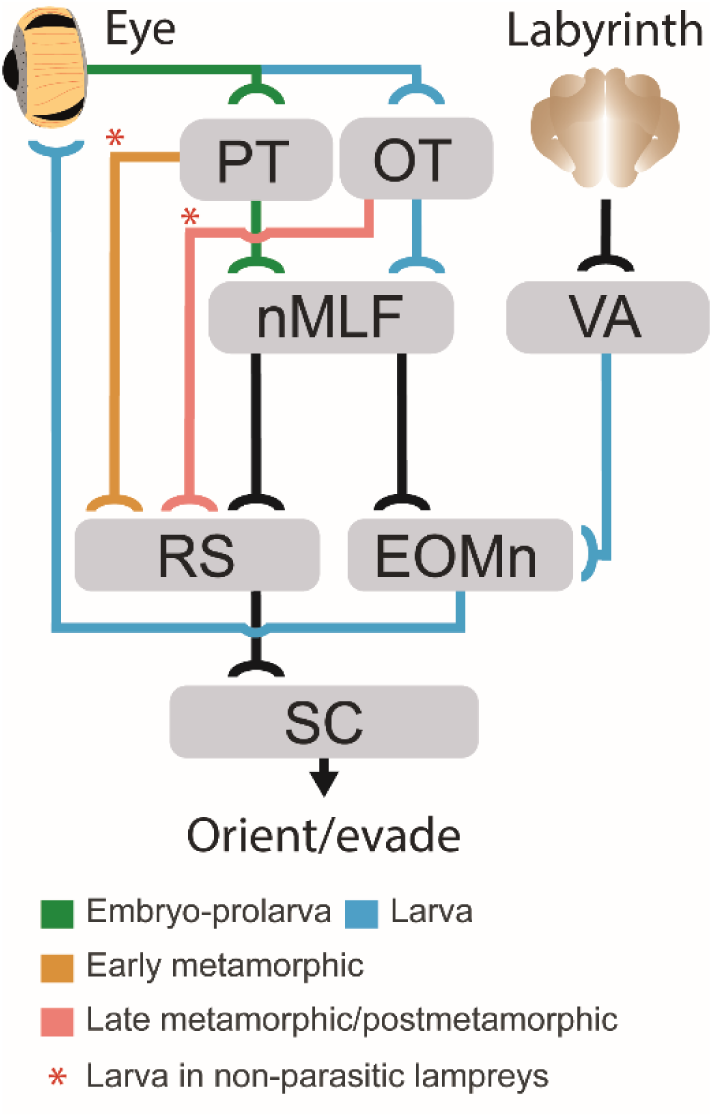
Development of gaze-controlling circuits in lampreys. Schematic showing the appearance stage of the main subcortical pathways mediating gaze control during development. Pathways that develop during embryo-prolarva stage are indicated in green, those that are stablished during the larval period in blue, those that appear at the beginning of metamorphosis in orange, and those that develop only in late metamorphosis are indicated in red. The red asterisks denote pathways that in non-parasitic lampreys start developing during the larval stage. Abbreviations: PT Pretectum, OT Optic Tectum, MRRN Middle Rhombencephalic Reticulospinal Nucleus, nMLF Nucleus of the Medial Longitudinal Fasciculus, VA Vestibular Area, RS Reticulospinal Neurons, EOMn Extraocular Muscles Motor Nuclei, SC Spinal Cord.

Reconstructing the evolutionary history of the visual system has been challenging because of the scarce data available in both extant and fossil early vertebrates (Fritzsch, 1991). Thus, understanding its development in lampreys provides valuable insight given their key phylogenetic position (Sugahara et al., 2017). Recent work has suggested that ancient lampreys lacked a filter-feeding stage and larvae had features of modern adult lampreys, including well-developed eyes (Miyashita et al., 2021). While the appearance of new mechanisms was necessary to fulfil the new feeding style (Youson, 1981), the strong similarities of the lamprey visual system and the emergence order of its underlying circuits with that of other vertebrates suggests that its stepwise development is merely a developmental slowing down spanning several years of the visual system already stablished in ancient lampreys, and that is largely conserved in other vertebrates. This stepwise construction of the visual system offers a unique window to study the role and functioning of the different visuomotor circuits.

## Materials and methods

### Animals

Experiments were performed on 51/21 sea lampreys/Nothern Japanese brook lampreys (*Petromyzon marinus*/*Lethenteron* sp. N, a cryptic species of *L. reissneri*; Yamazaki and Goto, 1998; Yamazaki et al., 2006), 40/12 larvae, 4/0 metamorphic (transformers), 5/9 postmetamorphic, and 2/0 adult animals. The experimental procedures were approved by the *Xunta de Galicia* under the supervision of the University of Vigo Committee for Animal use in Laboratory in accordance with the current regulations of the European Union (directive 2010/63/EU) and the Spanish regulation (*Real Decreto* 53/2013). A part of this study (using *Lethenteron* sp. N) was performed in accordance with the Regulations on Animal Experimentation at University of Tsukuba. Specific approval is not needed for experimentation on fish under Japanese law, Act on Welfare and Management of Animals. Also, every effort was made to minimize suffering and reduce the number of animals used in the study. The larvae were captured in a tributary of the Miño river (Furnia river: 41°59’33.0 “N 8°41’33.0” W), the transformers were kindly supplied by local fishermen with the permissions from *Xunta de Galicia* and *Comandancia Naval del Miño-Capitanía do Porto de Camiña/Comandancia local da Policía Marítima de Portugal*, whereas adults were obtained from authorized commercial distributors. Larvae and postmetamorphic of *Lethenteron* sp. N were captured from the Kamo River, which flows through the middle of the Shougawa River, Toyama, Japan. Animals were kept in aquaria with an enriched environment and continuously aerated and filtered water.

### *Ex-vivo* preparation

To expose the eyes and analyze their movements in response to electric stimulation of the brain, mechanical stimulation of the labyrinth, and light stimulation of the eyes, as well as to perform electrophysiological recordings in different brain regions, an *ex-vivo* preparation was used. For this, specimens were deeply anesthetized with tricaine methanesulfonate dissolved in water (0.1 %, MS-222, Sigma). When they stopped responding to tactile stimuli, the head was dissected at the level of the fourth gill and immersed in ice-cooled artificial cerebrospinal fluid (aCSF), with the following composition (in mM): 125 NaCl, 2.5 KCl, 2 CaCl_2_, 1 MgCl_2_, 10 glucose and 25 NaHCO_3_, saturated with 95 % (vol/vol) O_2_/5 % CO_2_. The skin, muscles and dorsal cartilage of the head were quickly removed to expose the brain. The choroid plexus and pineal gland were also removed. Subsequently, the lateral skin was removed to expose the eyes and otic capsules (containing the vestibular organs). The viscera and all muscles were removed to avoid movement of the preparation. The preparation was allowed to recover for at least 30 min and always kept immersed in cold aCSF during the experiments.

To mechanically stimulate the labyrinth, the cartilage of the otic capsule was cut carefully, so that the labyrinth was not damaged, opening a small window. A thin glass capillary, mounted on a micromanipulator (model M-3333, Narishige), was then gently pushed against the labyrinth membranes for the stimulation. To lesion the nMLF, an incision was made immediately caudal to this nucleus using a microscalpel through the third ventricle, thus cutting its downstream projections to the brainstem.

### Eye tracking

To test whether eye movements were evoked in response to electric stimulation of different brain areas, and light or vestibular stimulation, a video camera (AmScope MD35) attached to a stereo microscope (Leica M60) was used to record the eyes. Analysis of eye movements was performed with DeepLabCut (Mathis et al., 2018), a Python-based software package that uses artificial neural networks to estimate poses. To analyze the videos, one to four labels were first placed on each filmed eye in 20 key frames, and then training of the neural network was performed. After evaluating the training, the trained network was used to extract the position of the eyes throughout the video of interest. When several labels were used (in larger eyes), the trajectories of the eye labels were averaged to minimize errors. In some cases, a label was placed on the preparation and used to subtract small movements that might occur in the preparation due to muscle remnants in response to stimulation, or due to camera movements.

### Anatomical tract tracing

To study neuronal connections, tracer injections were performed in the AON, PT, retina, MRRN and SC. Injections were performed with glass micropipettes (borosilicate; od = 1.5 mm, id = 1.17 mm; Hilgenberg) with a tip diameter of 10-20 μm or sharpened tungsten pins. The glass micropipettes were attached to a holder connected to a pressurized air supply and this holder was placed on a micromanipulator (model M-3333, Narishige). Using the pressurized air system, the contents of the micropipette (tracer), which consisted of 50-200 nL of 20 % (wt/vol) Neurobiotin (Vector Laboratories) in aCSF containing Fast Green (Vector Laboratories), were injected into the brain (in the AON, PT or MRRN). Injections were also performed using dextran amine-tetramethylrhodamine (3 kDa; Molecular Probes). Alternatively, sharpened tungsten pins were mounted on a shaft and fluorescent dextran-amine (tetramethylrhodamine, 3,000 Da, Invitrogen, D3308; Alexa Fluor 488, 10,000 Da, Invitrogen, D22910) was recrystallized onto the tip of the pins (according to Glover, 1995). The crystal was inserted in the brain, retina or SC allowing anterograde or retrograde tracing. After injection, the brains were immersed in aCSF at 4 °C for 24 to 48 h in darkness to allow tracer transport, fixed in 4 % formaldehyde and 14 % saturated picric acid in 0.1 M saline phosphate buffer (PBS), pH 7.4, for 12-24 h, and cryoprotected in 20 % (wt/vol) sucrose in PBS for 3-12 h. Subsequently, they were embedded in OCT compound (Tissue-Tek, Sakura) and transverse sections of 30 μm thickness were cut in a cryostat (Leica cm1950) and collected on gelatinized slides. Before sectioning, some samples were dehydrated and clarified with BABB (1:2 mixture of benzyl alcohol and benzyl benzoate).

To detect the Neurobiotin tracer, sections were incubated in Cy2-conjugated streptavidin (Jackson ImmunoResearch) 1:1000 in blocking solution (1 % bovine serum albumin, 10 % sheep serum, 0.1 % sodium azide and 0.3 % Triton X-100 in PBS). A Nissl stain (Molecular Probes) was also added (1:500). Incubation was performed in a dark humid chamber for 2 h, and after three 10 min washes with PBS they were mounted in glycerol (Panreac).

### Electrophysiology

Extracellular recordings were performed in reticulospinal neurons in response to electrical stimulation of the AON, PT and OT, and to mechanical stimulation of the labyrinth using custom tungsten microelectrodes (∼1-5 MΩ). EMG recordings were also performed in the extraocular muscles. The recording electrodes were connected to an amplifier (AC differential amplifier, model 1700, AM systems) and the signal obtained digitized at 20 kHz using pClamp 10.4. The electrodes were operated using a micromanipulator (model M-3333, Narishige) that allowed their stable and precise positioning.

For electrical stimulation, borosilicate pipettes filled with aCSF were used, connected to a stimulus isolation unit (MI401). The stimulation intensity was set according to the threshold strength (usually 10–100 μA) necessary to evoke neuronal activity. For mechanical stimulation, a small window was made into the otic capsule, from which the vestibular system was directly stimulated by pressing the labyrinth with the tip of a pipette moved using a micromanipulator (model M-3333, Narishige).

### Image analysis

Photomicrographs were taken with a digital camera (Nikon DS-Ri2) mounted on a Nikon ECLIPSE Ni-E fluorescence microscope. Illustrations were made using Adobe Illustrator CC 2019, GIMP 2.1 (GNU image manipulator program), and Paint 3D. Images were only adjusted for brightness and contrast. The samples clarified with BABB were examined using a confocal laser micro-scope (LSM 510, Zeiss, Goettingen, Germany).

### Quantification and statistical analysis

For both electrophysiological and eye tracking data, analyses were performed using custom written functions in Matlab. For video recordings, eye positions were extracted with DeepLabCut (Mathis et al., 2018; see above). When only one eye was recorded frontally, to calculate the amplitude of the eye movement, the recorded eyes were extracted and measured in millimeters (mm) after the experiments, and the diameter of the eye was compared to its image so that a reliable conversion index from pixels to mm could be established. The video recorded eye movement amplitudes were consequently translated into mm. Using the diameter of the eye in the Z axis (distance from the optic nerve entrance to the cornea), its angular displacement could be calculated using trigonometric functions. To analyze the degree of eye coordination, a linear regression analysis was performed by calculating Pearson’s coherence coefficient.

To compare the signals evoked after each of the electric stimulation pulses, the integral under the curves were calculated after full rectification of the signals using trapezoidal numerical integration (‘trapz’ function) and the resulted values were normalized to the first pulse. To investigate the polysynaptic nature of the extracellular responses recorded, onset latencies were extracted by calculating the time from stimulus application to the first spike of the neuronal response. To compare latency times among different developmental stages, and the differences in MRRN activity in response to OT stimulation before and after nMLF lesioning, unpaired *t*-tests were used. The number of animals (N) and the number of experiments performed (n) are indicated where applicable. The degree of significance is indicated as follows: *P < 0.05, **P < 0.01, ***P < 0.001.

## Acknowledgements

We thank Professor Sten Grillner for his constant support, Eduardo Pena for hardware contribution, and Dr. Yuji Yamazaki for contributing resources.

## Author contributions

M.B, C.G.-N, D.G.S., M.A.P., and J.P.-F. conceived the study and designed the experiments. M.B, C.G.-N, D.G.S., C.J.-L., and J.P.-F performed the experiments. M.B, C.G.-N, D.G.S., and J.P.-F analyzed the data and designed the figures. M.B and J.P.-F wrote the manuscript with help from D.G.S. and M.A.P., with input from all the authors. J.P.-F supervised and funded the study.

## Data availability

All data are available on request.

## Supplementary Figures

**Fig. S1.**
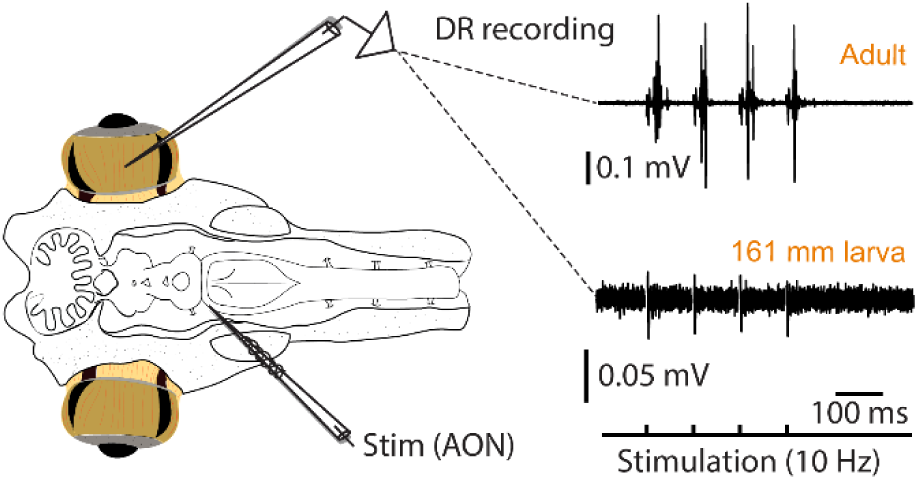
Extraocular muscles show similar activity in both larvae and adult lampreys. EMG activity in the dorsal rectus (DR) in of an adult lamprey (top trace) and a 161 mm larva (bottom trace) in response to a four pulses stimulation (10 Hz) of the anterior octavomotor nucleus (AON).

**Fig. S2.**
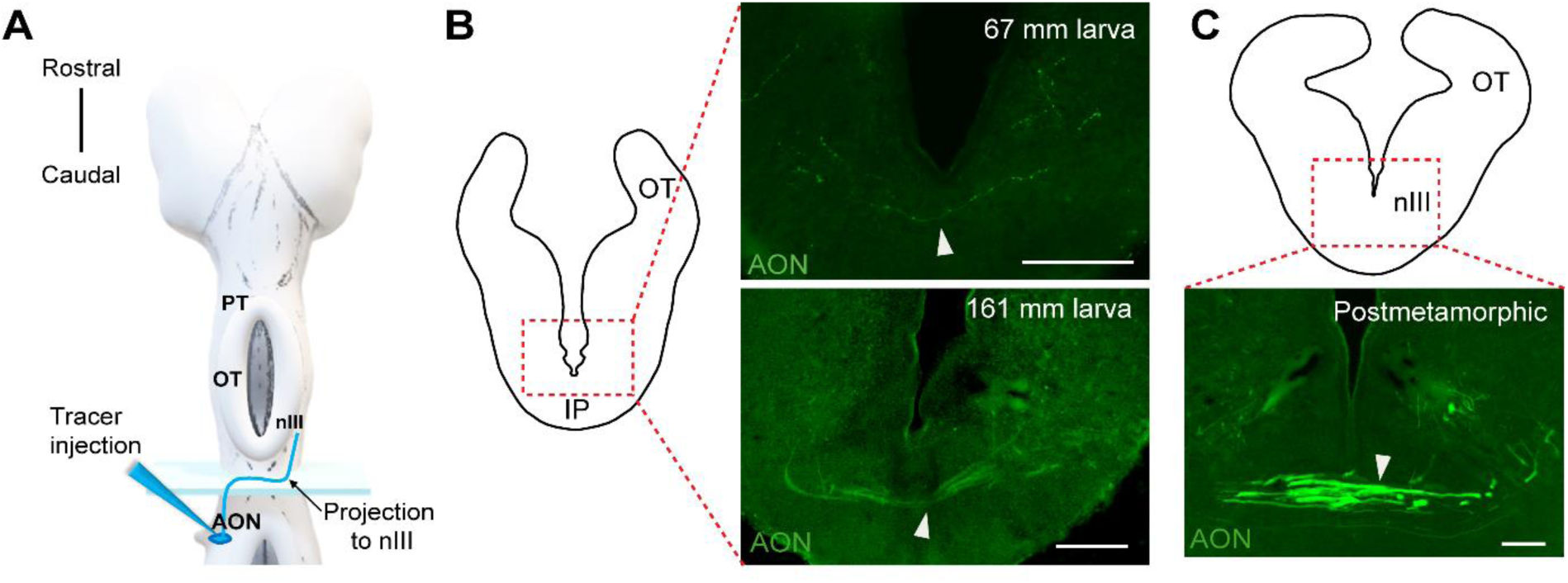
AON fibers crossing to the oculomotor nucleus. Schematic dorsal view of the lamprey larva brain indicating the tracer injection site in the anterior octavomotor nucleus (AON) from where the projections (blue line) that reach the oculomotor nucleus (nIII) originate. The rectangle indicates the location of the photomicrographs in B and C. (B) Anterogradely labeled fibers crossing at the level of the interpeduncular nucleus (IP) in a 67 mm larva (top) and a 161 mm larva (bottom). (C) Fibers crossing at the same location than B in a postmetamorphic animal. Scale bar = 100 µm B and C.

**Fig. S3.**
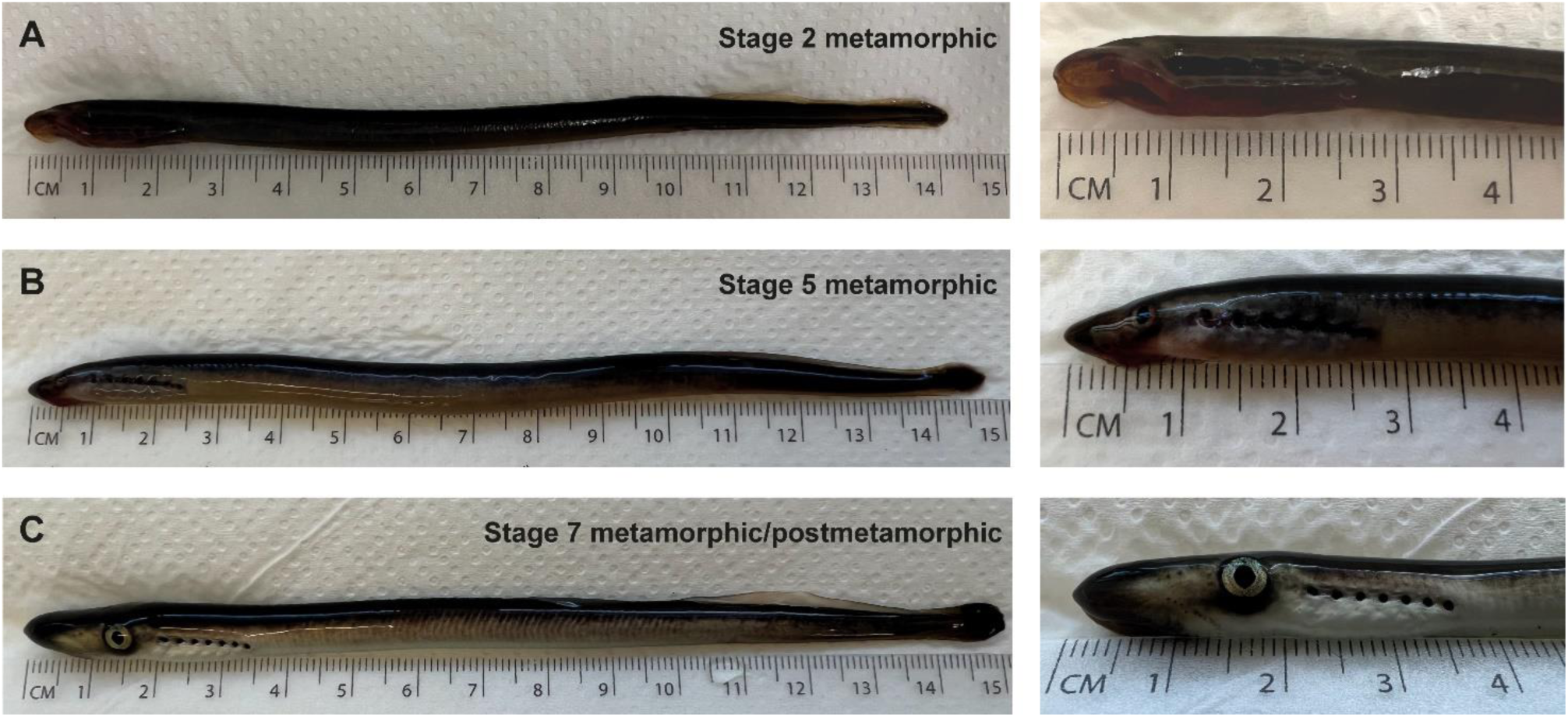
Representative metamorphic animals. (A-C) Photographs of representative metamorphic animals used in this study. A stage 2 metamorphic animal is shown in (A), a stage 5 in (B), and a stage7/postmetamorphic animal is shown in (C). A more detailed view of the head is shown in the right. The stage classification is based on Youson and Potter (1979).

## References

1. Branoner, F., Chagnaud, B. P. and Straka, H. (2016). Ontogenetic Development of Vestibulo-Ocular Reflexes in Amphibians. Front. Neural Circuits 10, 91.

2. Capantini, L., von Twickel, A., Robertson, B. and Grillner, S. (2017). The Pretectal Connectome in Lamprey. J. Comp. Neurol. 525, 753–772.

3. Cornide-Petronio, M. E., Barreiro-Iglesias, A., Anadón, R. and Rodicio, M. C. (2011). Retinotopy of Visual Projections to the Optic Tectum and Pretectum in Larval Sea Lamprey. Exp. Eye Res. 92, 274–281.

4. Dayton, G. O. J., Jones, M. H., Aiu, P., Rawson, R. A., Steele, B. and Rose, M. (1964). Developmental Study of Coordinated Eye Movements in the Human Infant. I. Visual Acuity in the Newborn Human: A Study Based on Induced Optokinetic Nystagmus Recorded by Electro-Oculography. Arch. Ophthalmol. 71, 865–870.

5. de Arriba, M del C. and Pombal, M. A. (2007). Afferent Connections of the Optic Tectum in Lampreys: An Experimental Study. Brain Behav. Evol. 69, 37–68.

6. de Miguel, E., Rodicio, M. C. and Anadón, R. (1990). Organization of the Visual System in Larval Lampreys: An HRP Study. J. Comp. Neurol. 302, 529–542.

7. Dickson, D. H. and Collard, T. R. (1979). Retinal Development in the Lamprey (*Petromyzon marinus* L.): Premetamorphic Ammocoete Eye. Am. J. Anat. 154, 321–336.

8. Donovan, T., Dunn, K., Penman, A., Young, R. J. and Reid, V. M. (2020). Fetal Eye Movements in Response to a Visual Stimulus. Brain Behav. 10, e01676.

9. du Lac, S. and Knudsen, E. I. (1991). Early Visual Deprivation Results in a Degraded Motor Map in the Optic Tectum of Barn Owls. Proc. Natl. Acad. Sci. U. S. A. 88, 3426–3430.

10. Easter, S. S. J. and Nicola, G. N. (1997). The Development of Eye Movements in the Zebrafish (*Danio rerio*). Dev. Psychobiol. 31, 267–276.

11. El Manira, A., Pombal, M. A. and Grillner, S. (1997). Diencephalic Projection to Reticulospinal Neurons Involved in the Initiation of Locomotion in Adult Lampreys *Lampetra fluviatilis*. J. Comp. Neurol. 389, 603–616.

12. Forsthofer, M. and Straka, H. (2023). Homeostatic Plasticity of Eye Movement Performance in Xenopus Tadpoles Following Prolonged Visual Image Motion Stimulation. J. Neurol. 270, 57–70.

13. Fritzsch, B. (1991). Ontogenetic Clues to the Phylogeny of the Visual System. In The Changing Visual System: Maturation and Aging in the Central Nervous System (ed. P. Bagnoli and W. Hodos), pp. 33–49. Boston, MA: Springer US.

14. Fritzsch, B. (1998). Evolution of the Vestibulo-Ocular System. Otolaryngol. Head Neck Surg. 119, 182–192.

15. Fritzsch, B., Sonntag, R., Dubuc, R., Ohta, Y. and Grillner, S. (1990). Organization of the Six Motor Nuclei Innervating the Ocular Muscles in Lamprey. J. Comp. Neurol. 294, 491–506.

16. Giolli, R. A., Blanks, R. H. and Lui, F. (2006). The Accessory Optic System: Basic Organization with an Update on Connectivity, Neurochemistry, and Function. Prog. Brain Res. 151, 407–440.

17. Glover, J. (1995). Retrograde and Anterograde Axonal Tracing with Fluorescent Dextran-Amines in the Embryonic Nervous System. Neurosci. Prot. 30, 1–13.

18. Grillner, S. and El Manira, A. (2020). Current Principles of Motor Control, with Special Reference to Vertebrate Locomotion. Physiol. Rev. 100, 271–320.

19. Hardisty, M. W. (2006). Lampreys: Life without Jaws. Cardigan: Forrest Text.

20. Horn, E., Lang, H. G. and Rayer, B. (1986). The Development of the Static Vestibulo-Ocular Reflex in the Southern Clawed Toad, Xenopus laevis. I. Intact Animals. J. Comp. Physiol. A. 159, 869–878.

21. Huang, Y. and Neuhauss, S. C. F. (2008). The Optokinetic Response in Zebrafish and its Applications. Front. Biosci. 13, 1899–1916.

22. Jones, M. R., Grillner, S. and Robertson, B. (2009). Selective Projection Patterns from Subtypes of Retinal Ganglion Cells to Tectum and Pretectum: Distribution and Relation to Behavior. J. Comp. Neurol. 517, 257–275.

23. Kardamakis, A. A., Pérez-Fernández, J. and Grillner, S. (2016). Spatiotemporal Interplay between Multisensory Excitation and Recruited Inhibition in the Lamprey Optic Tectum. Elife 5, 10.7554/eLife.16472.

24. Kardamakis, A. A., Pérez-Fernández, J. and Grillner, S. (2017). Networks in the Lamprey Optic Tectum. In Handbook of Brain Microcircuits (eds. G. M. Shepherd and S. Grillner), pp. 467–474. Oxford: Oxford University Press.

25. Kardamakis, A. A., Saitoh, K. and Grillner, S. (2015). Tectal Microcircuit Generating Visual Selection Commands on Gaze-Controlling Neurons. Proc. Natl. Acad. Sci. U. S. A. 112, E1956–65.

26. Kennedy, M. C. and Rubinson, K. (1977). Retinal Projections in Larval, Transforming and Adult Sea Lamprey, Petromyzon marinus. J. Comp. Neurol. 171, 465–479.

27. Kleerekoper, H. (1972). The Sense Organs. In Biology of Lampreys, Vol. 2 (eds. M. W. Hardisty and I. C. Potter), pp. 373–404. London: Academic Press.

28. Land, M. F. (2015). Eye Movements of Vertebrates and their Relation to Eye Form and Function. J. Comp. Physiol. A. Neuroethol Sens. Neural Behav. Physiol. 201, 195–214.

29. Masseck, O. A. and Hoffmann, K. P. (2009). Comparative Neurobiology of the Optokinetic Reflex. Ann. N. Y. Acad. Sci. 1164, 430–439.

30. Mathis, A., Mamidanna, P., Cury, K. M., Abe, T., Murthy, V. N., Mathis, M. W. and Bethge, M. (2018). DeepLabCut: Markerless Pose Estimation of User-Defined Body Parts with Deep Learning. Nat. Neurosci. 21, 1281–1289.

31. Miyashita, T., Gess, R. W., Tietjen, K. and Coates, M. I. (2021). Non-Ammocoete Larvae of Palaeozoic Stem Lampreys. Nature 591, 408–412.

32. Ocaña, F. M., Suryanarayana, S. M., Saitoh, K., Kardamakis, A. A., Capantini, L., Robertson, B. and Grillner, S. (2015). The Lamprey Pallium Provides a Blueprint of the Mammalian Motor Projections from Cortex. Curr. Biol. 25, 413–423.

33. Pérez-Fernández, J., Barandela, M. and Jiménez-López, C. (2021). The Dopaminergic Control of Movement-Evolutionary Considerations. Int. J. Mol. Sci. 22, 11284. doi: 10.3390/ijms222011284.

34. Pérez-Fernández, J., Kardamakis, A. A., Suzuki, D. G., Robertson, B. and Grillner, S. (2017). Direct Dopaminergic Projections from the SNc Modulate Visuomotor Transformation in the Lamprey Tectum. Neuron 96, 910–924.e5.

35. Pérez-Fernández, J., Stephenson-Jones, M., Suryanarayana, S. M., Robertson, B. and Grillner, S. (2014). Evolutionarily Conserved Organization of the Dopaminergic System in Lamprey: SNc/VTA Afferent and Efferent Connectivity and D2 Receptor Expression. J. Comp. Neurol. 522, 3775–3794.

36. Pombal, M. A., Rodicio, M. C. and Anadón, R. (1994). Development and Organization of the Ocular Motor Nuclei in the Larval Sea Lamprey, *Petromyzon marinus* L.: An HRP Study. J. Comp. Neurol. 341, 393–406.

37. Pombal, M. A., Rodicio, M. C. and Anadón, R. (1996). Secondary Vestibulo-Oculomotor Projections in Larval Sea Lamprey: Anterior Octavomotor Nucleus. J. Comp. Neurol. 372, 568–580.

38. Rovainen, C. M. (1976). Vestibulo-Ocular Reflexes in the Adult Sea Lamprey. J. Comp. Physiol. 112, 159–164.

39. Saitoh, K., Menard, A. and Grillner, S. (2007). Tectal Control of Locomotion, Steering, and Eye Movements in Lamprey. J. Neurophysiol. 97, 3093–3108.

40. Stephenson-Jones, M., Samuelsson, E., Ericsson, J., Robertson, B. and Grillner, S. (2011). Evolutionary Conservation of the Basal Ganglia as a Common Vertebrate Mechanism for Action Selection. Curr. Biol. 21, 1081–1091.

41. Straka, H. and Baker, R. (2013). Vestibular Blueprint in Early Vertebrates. Front. Neural Circuits 7, 182.

42. Sugahara, F., Murakami, Y., Pascual-Anaya, J. and Kuratani, S. (2017). Reconstructing the Ancestral Vertebrate Brain. Dev. Growth Differ. 59, 163–174.

43. Suryanarayana, S. M., Pérez-Fernández, J., Robertson, B. and Grillner, S. (2021). The Lamprey Forebrain – Evolutionary Implications. Brain Behav. Evol., 1-16.

44. Suryanarayana, S. M., Pérez-Fernández, J., Robertson, B. and Grillner, S. (2020). The Evolutionary Origin of Visual and Somatosensory Representation in the Vertebrate Pallium. *Nat*. Ecol. Evol. 4, 639–651.

45. Suzuki, D. G., Fukumoto, Y., Yoshimura, M., Yamazaki, Y., Kosaka, J., Kuratani, S. and Wada, H. (2016). Comparative Morphology and Development of Extra-Ocular Muscles in the Lamprey and Gnathostomes Reveal the Ancestral State and Developmental Patterns of the Vertebrate Head. Zoological Lett. 2, 10-016-0046-3. eCollection 2016.

46. Suzuki, D. G. and Grillner, S. (2018). The Stepwise Development of the Lamprey Visual System and its Evolutionary Implications. Biol. Rev. Camb. Philos. Soc. 93, 1461–1477.

47. Suzuki, D. G., Murakami, Y., Escriva, H. and Wada, H. (2015a). A Comparative Examination of Neural Circuit and Brain Patterning between the Lamprey and Amphioxus Reveals the Evolutionary Origin of the Vertebrate Visual Center. J. Comp. Neurol. 523, 251–261.

48. Suzuki, D. G., Murakami, Y., Yamazaki, Y. and Wada, H. (2015b). Expression Patterns of Eph Genes in the “Dual Visual Development” of the Lamprey and their Significance in the Evolution of Vision in Vertebrates. Evol. Dev. 17, 139–147.

49. Suzuki, D. G., Pérez-Fernández, J., Wibble, T., Kardamakis, A. A. and Grillner, S. (2019). The Role of the Optic Tectum for Visually Evoked Orienting and Evasive Movements. Proc. Natl. Acad. Sci. U. S. A. 116, 15272–15281.

50. Ullén, F., Deliagina, T. G., Orlovsky, G. N. and Grillner, S. (1997). Visual Pathways for Postural Control and Negative Phototaxis in Lamprey. J. Neurophysiol. 78, 960–976.

51. Villar-Cerviño, V., Abalo, X. M., Villar-Cheda, B., Meléndez-Ferro, M., Pérez-Costas, E., Holstein, G. R., Martinelli, G. P., Rodicio, M. C. and Anadón, R. (2006). Presence of Glutamate, Glycine, and Gamma-Aminobutyric Acid in the Retina of the Larval Sea Lamprey: Comparative Immunohistochemical Study of Classical Neurotransmitters in Larval and Postmetamorphic Retinas. J. Comp. Neurol. 499, 810–827.

52. von Twickel, A., Kowatschew, D., Salturk, M., Schauer, M., Robertson, B., Korsching, S., Walkowiak, W., Grillner, S. and Pérez-Fernández, J. (2019). Individual Dopaminergic Neurons of Lamprey SNc/VTA Project to both the Striatum and Optic Tectum but Restrict Co-Release of Glutamate to Striatum Only. Curr. Biol. 29, 677–685.e6.

53. Walls, G. L. (1962). The Evolutionary History of Eye Movements. Vision Res. 2, 69–80.

54. Wang, L., Liu, M., Segraves, M. A. and Cang, J. (2015). Visual Experience is Required for the Development of Eye Movement Maps in the Mouse Superior Colliculus. J. Neurosci. 35, 12281–12286.

55. Wibble, T., Pansell, T., Grillner, S. and Pérez-Fernández, J. (2022). Conserved Subcortical Processing in Visuo-Vestibular Gaze Control. Nat. Commun. 13, 4699-022-32379-w.

56. Yamazaki, Y. and Goto, A. (1998). Genetic Structure and Differentiation of Four *Lethenteron* Taxa from the Far East, Deduced from Allozyme Analysis. Environ. Biol. Fishes 52, 149–161.

57. Yamazaki, Y., Yokoyama, R., Nishida, M. and Goto, A. (2006). Taxonomy and Molecular Phylogeny of *Lethenteron* Lampreys in Eastern Eurasia. J. Fish Biol. 68, 251–269.

58. Youson, J. H. (1981). The Alimentary Canal. In Biology of Lampreys, Vol. 3 (eds. M. W. Hardisty and I. C. Potter), pp. 95–189. London: Academic Press.

59. Youson, J. H. and Potter, I. C. (1979). A Description of the Stages in the Metamorphosis of the Anadromous Sea Lamprey, *Petromyzon marinus* L. Can. J. Zool. 57, 1808–1817.

60. Zompa, I. C. and Dubuc, R. (1996). A Mesencephalic Relay for Visual Inputs to Reticulospinal Neurones in Lampreys. Brain Res. 718, 221–227.

61. Zompa, I. C. and Dubuc, R. (1998). Diencephalic and Mesencephalic Projections to Rhombencephalic Reticular Nuclei in Lampreys. Brain Res. 802, 27–54.

